# Changes in the cellular composition of the endometrium during the implantation window are associated with recurrent pregnancy loss

**DOI:** 10.1101/2024.10.22.619501

**Authors:** Ian G. Reddin, Jennifer E. Pearson-Farr, Anna H. Turaj, Sean H. Lim, Rohan M. Lewis, Ying C Cheong, Jane K. Cleal

**Affiliations:** School of Cancer Sciences, University of Southampton Faculty of Medicine, Southampton, SO16 6YD, UK; Bio-R Bioinformatics Research Facility, University of Southampton Faculty of Medicine, Southampton, SO16 6YD, UK; Human Development and Health, University of Southampton, Faculty of Medicine, Tremona Road, Southampton, SO16 6YD, UK; University Hospital Southampton NHS Foundation Trust, Coxford Road, Southampton, SO16 5YA, UK; Institute for Life Sciences, University of Southampton, Highfield, Southampton SO17 1BJ, UK

## Abstract

Recurrent pregnancy loss (RPL) affects 1-2% of women trying to conceive, yet in many cases the causes remain unclear. Endometrial function is central to the establishment and maintenance of pregnancy, and endometrial dysfunction may underlie RPL. A greater understanding of the endometrial cell populations and their interactions in women with and without RPL may identify markers of endometrial receptivity and the likelihood of pregnancy success. Single cell RNA sequencing was performed on RPL (n = 3) and control (n = 4) endometrial biopsies collected at days 21-24 of the menstrual cycle, the window of implantation. 10,022 cells were clustered and nine major individual cell types were characterised. Further analysis identified six distinct endometrial stromal cell (EnSCs) and three natural killer (NK) cell sub-populations. In RPL, there were changes in the abundance of specific endometrial stromal and NK cell subpopulations with associated differences in cellular communication between the cell types related to the Wnt pathway and angiogenesis. This is consistent with NK cell signalling orchestrating the difference in abundance of stromal cells and regulating processes needed for successful implantation. These changes in RPL endometrium provide further evidence for an endometrial cause of RPL and identify specific mechanisms for future study.

## Introduction

Recurrent pregnancy loss (RPL), the loss of three or more clinical pregnancies [1], affects 1-2% of women trying to conceive yet for many cases the causes are unknown [2]. Each successive pregnancy loss increases the risk of subsequent pregnancy loss, which further increases with maternal age, implicating uterine factors in RPL [3]. Altered endometrial receptivity to the embryo during the window of implantation (days 21-24 of the menstrual cycle) and reduced endometrial gland secretions may contribute to RPL [4].

Potential biomarkers of receptive endometrium have been proposed [5–7], however a consistent gene profile predictive of pregnancy outcome is not yet established [8]. This may reflect the heterogeneity of endometrium and cell specific expression patterns across the menstrual cycle. Single-cell gene expression has been used to identify distinct endometrial cell populations based on their transcriptional states at different stages of the menstrual cycle. However, it remains unclear whether these cell populations or their gene profiles are altered in cases of RPL [4, 9].

The endometrial cell populations include luminal epithelium and glandular epithelium with ciliated and secretory cells [4] that support implantation and successful pregnancy [10]. The underlying stroma contains fibroblast-like endometrial stromal cells (EnSCs), macrophages, dendritic cells (DCs), endothelium, perivascular (PV) cells, neutrophils and uterine natural killer (uNK) cells [9, 11]. Successful embryo implantation requires appropriate cell-to-cell communication with the discreet cell populations adopting a coordinated receptive state during the window of implantation. These adaptations include hormonal cues that stimulate EnSCs differentiation into specialised decidual stromal cells (DSCs), accompanied by an increase in uNK cells in the endometrium [19]. This provides the immunological and nutritional environment that is required for embryo implantation. Whether there are differences in these cell populations or their interactions in RPL during the window of implantation is not known.

This study uses single cell RNA sequencing of endometrium from the window of implantation in women with and without RPL to identify endometrial gene expression and cell population changes associated with RPL. Nine cell populations in the endometrium during the window of implantation were characterised. Six distinct endometrial stromal cell subpopulations (SC1-6) were identified, with probable functions ranging from extracellular matrix arrangement to angiogenesis and the innate immune response. An *ACTA2*^+^*TAGLN*^+^ EnSCs subpopulation (SC4) representing a myofibroblast phenotype was less abundant in RPL, while an *IGF1*^+^/*DIO2*^+^/*SCARA5*^-^ subpopulation (SC5) was more abundant in RPL vs control endometrium. These differences in EnSC cell populations in RPL may be orchestrated by changes to the three uNK cell subpopulations identified. Indeed, NK2 potentially stimulates differentiation of NK cells to NK1 and SC5 to SC4 via *AREG*, and both NK1 and NK2 were less abundant in RPL. NK3 has a role in immune cell recruitment via chemokines *CCL3/4/5* and *XCL1* and was more abundant in RPL.

In RPL endometrium compared to control, differences in cellular communication between the cell types related to the Wnt signalling pathway and angiogenesis. Regulation of these processes is vital for successful implantation and pregnancy, so taken together with differences in cell type abundance observed in our analysis, this contributes to the understanding of RPL and provides direction for future research into RPL.

## Results and discussion

### Single cell characterisation of RPL and control endometrium during the window of implantation

Single cell RNA sequencing (scRNAseq) identified cellular interactions during the window of implantation (day 21-24 of the menstrual cycle) in endometrium from RPL (loss of ≥3 pregnancies) and control participants who met the criteria for egg donation. Tissue biopsies digested into single-cell suspensions underwent scRNAseq (10x Chromium platform) to identify the specific endometrial cell populations and their molecular signatures. Following quality control (QC) for low-quality cells, the data for one control sample was removed and 9,250 cells were retained for further analysis; 5,051 for control (n = 4 participants) and 4,199 for RPL (n = 3 participants; **Supplementary Figure S1A-C**).

Unsupervised clustering (Seurat R package) of the combined dataset identified nine cell populations (**Figure 1A**). Known individual marker genes *GPX3* (epithelial cell), *CD14* (macrophage), *CD1A* (dendritic cell), *CD3D* (T-cell), *MCAM* (endothelial cell), *DCN* (stromal cell), *MS4A1* (B-cell), *IL2RB* (natural killer cell) and *TPSB2* (mast cell) were used to determine cell type (**Figure 1B, Supplementary Figure S1D**). Further marker genes and literary evidence were used to confirm the given identities for each cell type (**Figure 1C**).

**Figure 1:**
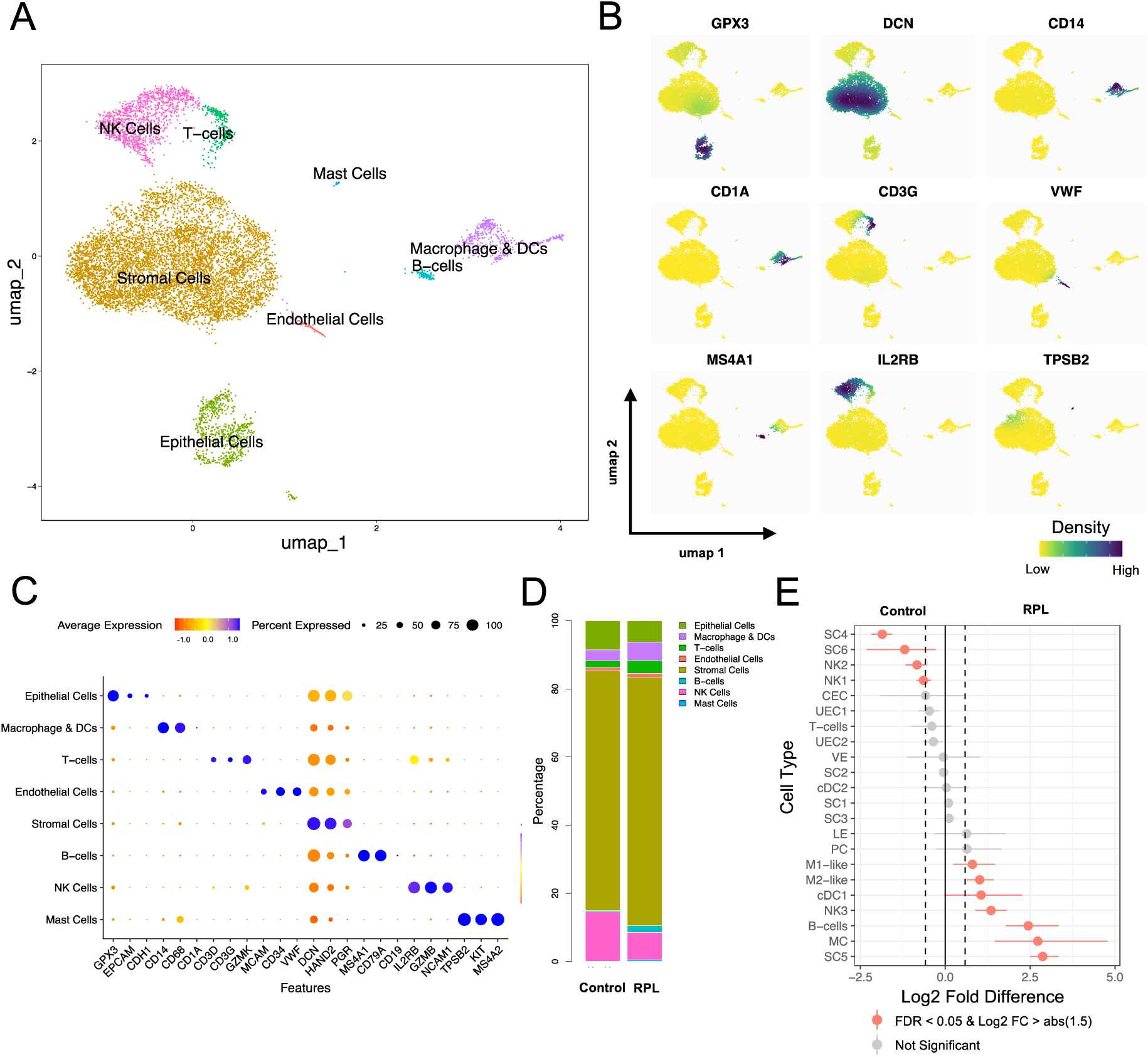
Single cell overview of the human endometrium during the secretory phase. **(A)** Uniform manifold approximation and projection (UMAP) dimensional reduction to visualise the 9,250 cells clustered into nine major cell types based initially on single gene markers for which expression density was visualised in individual UMAPS **(B)**. Further *a priori* knowledge and expression of additional marker genes as visualised in a dotplot **(C)** for all cells was used to confirm cell types. **(D)** Stacked bar chart showing cellular proportions in both experimental groups, and following further clustering of cell subtypes significant cell proportion differences between groups were identified **(E)**. SC1-6 = endometrial stromal cell 1-6, NK1-3 = Natural killer cell 1-3, CEC = ciliated epithelial cell, UEC1-2 = unciliated epithelial cell 1-2, VE = vascular endothelium, LE = lymphatic like endothelium, PC = pericytes, cDC1-2 = conventional dendritic cell 1-2, M1-like = M1-like macrophage, M2-like = M2-like macrophage, MC = mast cells.

All seven endometrial samples matched the gene expression profile in EnSCs and epithelial cells associated with progression of the menstrual cycle [4]. Genes highly expressed in mid/late-secretory phase EnSCs (*S100A4, DKK1, CRYAB, and FOXO1*) were the highest expressed in our EnSC population, while genes expressed highest in the late secretory phase (*IL15, FGF7*) were relatively low (**Supplementary Figure S1E**). Similarly, genes most expressed in epithelial cells in mid/late-secretory phase (metallothionein genes, *CXCL14*) were the highest expressed in our epithelial population (**Supplementary Figure S1F**). This indicates that EnSCs and epithelial cells, and therefore all endometrial tissue samples, were collected in the mid-secretory phase of the menstrual cycle.

### Differences in cell populations of RPL and control endometrium during the window of implantation

The endometrial stromal cell (EnSC) population was the most abundant in both the control (mean, 71.5%; range, 56.0-84.9%) and RPL (72.3%, 72.0-73.1%) groups, while uterine natural killer (uNK) cell (control, 14.0%, 8.4-19.5%; RPL, 9.9%, 9.0-10.1%) and epithelial cells (control, 8.3%, 1.4-14.4%; RPL, 6.0%, 5.7-7.5%) were second and third most abundant respectively (**Figure 1D**, **Supplementary Figure S1G, Supplementary Tables S1&S2**).

The nine endometrial cell types in the complete data set clustered further, resulting in 22 total endometrial cell subpopulations. Two EnSC populations (SC4 and SC6; fold difference = 3.6 and 2.3, FDR = 0.002 and 0.01 respectively) and two uNK cell subtypes (NK1 and NK2; fold difference = 1.56 and 1.78, FDR = 0.002) were significantly less abundant in RPL endometrium compared to control (**Figure 1E, Supplementary Table S3**).

Seven cell populations were proportionally more abundant in RPL endometrium: one EnSC subpopulation (SC5, fold difference = 7.3, FDR = 0.002) one uNK cell subpopulation (NK3, fold difference = 2.5, FDR = 0.002), two macrophage subtypes (M1-like, fold difference = 1.7, FDR = 0.009; M2-like, fold difference = 2, FDR = 0.002), one dendritic cell subpopulation (cDC1, fold difference = 2.1, FDR = 0.047), B-cells and mast cells (fold difference = 5.4 and 6.6 respectively, FDR = 0.002 for both), (**Figure 1E, Supplementary Table S3**).

Differences in cell populations in RPL compared to control endometrium during the window of implantation suggests altered restructuring of the endometrial stroma. The reduced presence of myofibroblast-like SC4 cells in RPL endometrium could either indicate a deficiency of this cell type in the samples or an incomplete or delayed differentiation from progenitor cells. In either scenario, the endometrial remodelling necessary for successful implantation may be impaired. Additionally, the RPL endometrium showed an enrichment of SC5 cells, which exhibit a senescent-like phenotype, suggesting an altered decidualization trajectory, which may relate to the altered uNK cell populations in RPL.

### Single cell RNA sequencing analysis of the endometrial stromal cell population identifies six subpopulations with potentially different functions

After QC a total of 6,501 EnSCs were available for further analysis. A total of 3,508 EnSCs from control endometrium and 2,993 from RPL endometrium were clustered individually. Cells in both groups clustered into four EnSC subtypes (**Supplementary Figure S2A&B**), three of which shared markers between control and RPL (control SC1, SC2 and SC3 with RPL SC1, SC2 and SC3 respectively), and one unique to each group (control, SC4 and RPL, SC5; **Supplementary Figure S2C&D**). The five total clusters were conserved when the cells from both groups were clustered together, and an additional EnSC subtype was identified when considering the complete data set (SC6; **Figure 2A, Supplementary Figure S2E&F**). The two clusters individually identified in control and RPL were enriched in the respective groups, 13.3% of control EnSCs were SC4 compared to only 3.6% of RPL EnSCs (FDR = 0.002), and 13.3% of RPL EnSCs were SC5 compared to only 1.6% of control EnSCs (FDR = 0.002). There was no significant difference in the remaining three subtypes observed in both cases (SC1: 34.6% in control vs 36% in RPL, FDR = 0.07; SC2: 31.9% vs 30%, FDR = 0.23; SC3: 17.6% vs 18.6%, FDR = 0.13), while SC6 was enriched in control compared to RPL (0.9% in control, 0.4% in RPL, FDR = 0.01; **Figure 1E&2B, Supplementary Figure S2G, Supplementary Table S3**). This pattern of EnSC phenotype abundance was largely followed in each individual sample. SC4 was more abundant than SC5 in 3 of the 4 control samples and SC5 more abundant than SC4 in 2 of the 3 RPL samples, while the remaining 4 EnSC subtypes were present in similar abundance across all samples (**Figure 2C**). In addition, a larger proportion of control cells were in the S-phase of the cell cycle compared to RPL cells (FDR = 0.001; **Figure 2D, Supplementary Figure S3A, Supplementary Table S4**).

**Figure 2:**
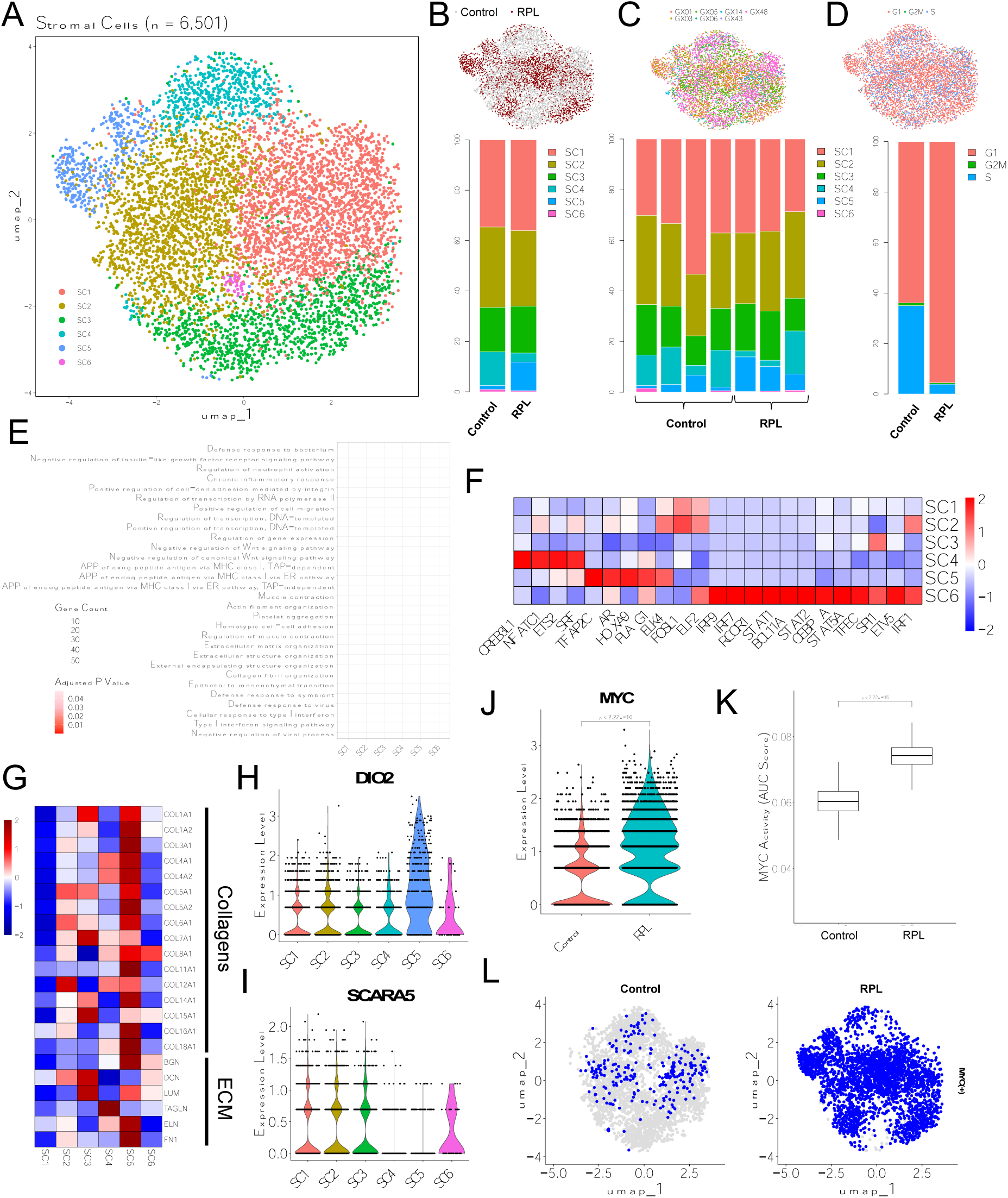
Characterisation of endometrial stromal cells (EnSCs) in control and RPL patient groups in the secretory phase. **A)** UMAP of 6,501 EnSCs were clustered into six individual subtypes. **B)** UMAP at top shows breakdown of EnSC by patient group, while stacked bar chart at bottom shows the relative abundance of EnSCs in each patient group. **C)** UMAP at top shows breakdown of EnSC by patient type, stacked bar charts at the bottom show relative proportion of each EnSC in individual patients. **D)** UMAP at top shows breakdown of cells by cell cycle phase, bar chart at bottom the relative proportion of cells that are in each cell cycle phase for both patient groups. **E)** Dotplot showing top 5 GOBP pathways associated with upregulated genes in each EnSC population. **F)** Differential transcription factor activity between the EnSC subtypes for transcription factors where there was a significant difference. **G)** Expression of collagen and extracellular matrix associated genes. The difference in expression between EnSCs shown in violin plots for H) *DIO2* and I) *SCARA5*, and between patient groups for J) *MYC* gene expression was different in control and RPL EnSCs (P < 2.22 x 10^-16^). **K)** Boxplot showing a significant difference in *MYC* transcription factor activity score between both patient groups (significance calculated using Wilcoxon rank sum test). **L)** UMAPS of *MYC* binary activity status in both patient groups, grey = OFF, blue = ON.

### A deficiency of myofibroblast-like SC4 in RPL endometrium at the window of implantation suggests delayed restructuring of the endometrial stroma

The top pathways associated with upregulated genes in SC4 included muscle contraction and actin filament organisation (**Figure 2E**) while *ACTA2* and *TAGLN* were the most significantly upregulated genes in control patient dominated SC4 (**Supplementary Table S5**). This suggests that SC4 may represent a myofibroblast phenotype. Transcription factors upregulated in SC4 include *CREB3L1* and *SRF* (**Figure 2F**), both of which are regulators of myofibroblast differentiation [12–14]. *ACTA2^+^* EnSC populations have been identified in endometrium [15] and in an integrated validation dataset as cluster 1 (**Supplementary Figure S3B&C**) [16]. Myofibroblasts perform functions that are vital in remodelling of the endometrium in the window of implantation, which include promotion of vascularisation and angiogenesis via the remodelling of endothelial cells. During a normal menstrual cycle *ACTA2^+^* EnSC differentiation is suspected to be reduced by factors in menstrual discharge which prevent scarring of the endometrium as it heals [17], however myofibroblasts/*ACTA2^+^* EnSCs have been identified in the proliferative phase of the menstrual cycle [15]. The low number of myofibroblast-like SC4 in RPL patients suggests that either this cell type is deficient in our RPL samples or they have not yet differentiated from a progenitor cell. In either case, restructuring of the endometrium for implantation could be incomplete.

### RPL endometrium were enriched with SC5 that resemble a senescent phenotype indicative of an altered decidualisation trajectory

The SC5 subpopulation of EnSCs was characterised by increased expression of collagen and extracellular matrix (ECM) genes identified by Qian et al (2020) [18] (**Figure 2G**). ECM-related pathways also dominated the top pathways based on genes upregulated in SC5 (**Figure 2E**). The gene *DIO2* was upregulated in SC5 (**Figure 2H, Supplementary Table S5**), while *SCARA5* was downregulated (**Figure 2I**), suggesting that this subpopulation of EnSCs may demonstrate characteristics of senescent EnSCs as described by Lucas *et al* (2020)[19]. An acute senescent cell phenotype is indicated by a permanent state of cell cycle arrest and the secretion of key molecules linked to biological processes that involve programmed tissue remodelling including inflammatory mediators [20]. Such processes are key to the initiation of decidualisation, whereby, prior to implantation, EnSCs differentiate into the specialised decidualised stromal cells (DSCs) to provide the environment required for embryo implantation. Indeed, this senescent cell phenotype has been linked to initiation of decidualisation [21]. *IGF1* was also upregulated in SC5 EnSCs (**Supplementary Table S5**) and has previously been identified as a marker gene for a stromal cell population (Rem-SC) that initiates decidualisation [21]. Together this suggests that our SC5 population is an *IGF1*^+^/*DIO2*^+^/*SCARA5*^-^ EnSC population that is important in the initiation of decidualisation. Intercellular interactions between EnSCs cells are upregulated upon decidualisation and vital for establishing a functional feto-maternal interface. This is accompanied by an increase in uNK cells, which may clear senescent cells. Appropriate cell-to-cell communication must balance the induction and clearance of senescent cells, allowing for an inflammatory response for successful embryo implantation; this may be dysregulated in RPL [19].

The differentiation pathway of the EnSC subpopulations during decidualisation can be modelled using trajectory analysis. Trajectory analysis using Monocle3 with SC5 as the root node indicated that SC2 and SC4 were next in the trajectory with similar pseudotime profiles (SC2 mean pseudotime score = 1.94, SC4 = 2.19), while SC1, SC3, and SC6 had higher pseudotime scores (**Supplementary Figure S3D, Supplementary Table S6**) indicating a subsequent induction. This suggests that SC4 and SC2 differentiate from SC5 as part of the stromal remodelling required for a receptive endometrium during the window of implantation. SC2 is likely the first decidualised EnSC subtype that differentiates from SC5 EnSCs. *IGF1R* and *ADAMTS5* are significantly upregulated in SC2 compared to other EnSC subtypes (**Supplementary Table S5**), resembling the decidualised *IGF1R^+^*/*ADAMTS5^high^* dRem-SC EnSC identified by Shi *et al* (2022) [22]. The top pathways associated with upregulated genes in SC2 were involved with regulation of gene expression (**Figure 2E**) which may coincide with an ability to differentiate into a number EnSCs with different phenotypes. The different transcriptional profiles of the EnSC subpopulations suggest they have differing roles (**Supplementary Figure S3E**).

### EnSC populations involved in endometrial homeostasis

EnSC populations SC1 and SC3 (not enriched in either group) and SC6 (enriched in control samples) have potential roles in maintaining endometrial homeostasis (**Figure 1E**). The top 5 gene ontology biological process (GOBP) pathways, based on differentially expressed genes in SC1 and SC6 included those involved in the immune response, and response to inflammation. The most upregulated (by fold change) gene in SC1 was chemokine *CXCL13* (**Supplementary Table S5**), which is expressed in endometrial epithelial cells during the secretory phase of the menstrual cycle and aberrantly expressed in endometriosis [23]. *PLA2G2A,* a gene that has been associated with an EnSC with high secretory ability (Sec-SC) [21] is also upregulated in SC1 (**Supplementary Table S5**).

Top upregulated genes in SC3 include *TIMP3*, which prevents degradation of ECM, and *SOX18*, which is involved in blood and lymphatic vessel development (**Supplementary Table S5**). This suggests that this EnSC subtype has a role in the remodelling of the endometrium during the secretory phase of the menstrual cycle. The top pathways associated with genes upregulated in SC3 EnSCs included regulation of the Wnt-signalling pathway, which is associated with endometrial structural and functional changes [24], and antigen processing and presentation (APP) indicating a role for SC3 in the endometrial immune response (**Figure 2E**). APP of exogenous antigen peptides via MHC class I suggests that this cell type presents peptides via cross-presentation and relies on endocytosis. SC3 EnSCs could exhibit phagocytotic activity via these APP pathways, and may contribute to ECM remodelling by endometrial cell clearance [25].

The small SC6 subpopulation resembles EnSC (*ISG15^+^* cluster) identified in the proliferative endometrium [15], and a small EnSCs cluster in the mid-secretory phase from an integrated dataset comprising two scRNA-seq datasets (**Supplementary Figure S3B&C**) [4, 26]. Genes associated with the interferon response including *MX1*, *IFIT1*, *ISG15*, *IFIT3*, and *MX2* are highly expressed in SC6 (**Supplementary Table S5**), which is reflected in pathway analysis where the top five pathways are interferon response or defence against viruses (**Figure 2E**). The transcription factors *STAT1*, *STAT2*, *IRF1*, *IRF7*, and *IRF9* were upregulated in SC6 (**Figure 2F**) supporting a role in regulating the immune system/response. The enrichment of SC6 in control endometrium suggest that the RPL endometrium is deficient in homeostatic regulation in response to external stimuli that should initiate an immune response.

### Differentially expressed genes in EnSCs may be associated with RPL

Several genes were differentially expressed in RPL EnSCs compared to control and when considering all cells regardless of cluster. While some differentially expressed genes were to be expected, particularly those expressed at higher levels in one of the clusters dominated by either patient group (e.g., *ACTA2* from SC4 upregulated in control, *DIO2* from SC5 upregulated in RPL), there were examples of genes being differentially expressed across all clusters. *DKK1*, *MTRNR2L8*, *CXCL13*, and *MYC* were notable genes upregulated in RPL endometrium, while *HLA-C* was higher in control endometrium (**Supplementary Table S7**).

*DKK1* is highly expressed in the mid-secretory phase of the menstrual cycle [27–29] and can inhibit the Wnt-signalling pathway, which mediates mesenchymal-epithelial interactions and regulates implantation in the murine endometrium [30]. Upregulation of *DKK1* in RPL endometrium may indicate dysregulation of the Wnt-signalling pathway in these individuals. *MTRNR2L8* is a pseudogene associated with anti-apoptotic roles and inflammation [31, 32], which are important for endometrial homeostasis [33, 34], while *CXCL13* is a chemo-attractant for B-cells [35], and associated with chronic endometritis/inflammation and implantation failure [36]. The roles associated with these genes suggest that they are important for maintaining a suitable environment for successful implantation/pregnancy and their dysregulation may increase the chances of pregnancy loss.

The transcription factor *MYC* has roles including cell cycle and apoptosis [37]. *MYC* gene expression in EnSCs is highest in the mid-secretory phase of the menstrual cycle, which is confirmed in the validation cohorts (**Supplementary Figure S3F**). In our cohort *MYC* was expressed at a higher level in EnSCs in RPL endometrium (fold change = 3.5, adjusted p-value < 0.0001, **Figure 2J**). Furthermore, the *MYC* transcription activity score (as calculated in SCENIC analysis) was significantly higher in RPL endometrium compared to control (**Figure 2K**) and based on a binary threshold for *MYC* activity (i.e. “ON” or “OFF”), MYC was considered “ON” in 2,766/2,993 (93%) of RPL EnSCs but “ON” in only 239/3,508 (7%) of control cells (**Figure 2L**). In mice overexpression of *Myc* has been identified as a compensatory mechanism for failure of EnSC decidualisation caused by *Mettl3* deficiency [38]. *METTL3* gene expression was not decreased in our RPL group. However, given our observation of a RPL dominated pre-decidualised EnSC subpopulation (SC5), there may have been a delay in decidualisation and the increased *MYC* expression is attempting to compensate for this delay. Altered expression of *MYC* has been associated with another endometrial condition, endometriosis [39], adding to our evidence that *MYC* expression in RPL endometrium should be studied further.

The *HLA-C* gene is upregulated in control endometrium (fold change = 3.4, p < 0.0001), and encodes a ligand of Killer cell immunoglobulin-like receptors (KIR) found on the surface of NK cells [40]. NK cells are associated with the clearance of senescent EnSCs via granule exocytosis [19]. Reduction in *HLA-C* expression in RPL EnSCs could result in a reduction in the clearance of senescent/senescent-like EnSCs like the RPL dominated SC5 EnSC population.

### Identification of cell subtypes in the immune microenvironment of endometrium

The immune cell subset consisted of 1,974 cells (1,084 control, 890 RPL) and reclustering resulted in the five previously identified immune cell populations (**Figure 3A**) characterised by marker genes for each cell type (**Figure 3B, Supplementary Table S8**). uNK cells were the most abundant immune cell type in both control (76.5%) and RPL (56.4%) endometrial samples but, considering all uNK cells, were not significantly different in abundance between the two groups (**Figure 3C, Supplementary Figure S4A**). The macrophage and dendritic cell population was the second most abundant immune cell type (15.4% control, 26.9% RPL) and was significantly enriched in RPL patients (proportion test, fold difference, 1.72, FDR = 0.002), as were B-cells (fold difference = 5.4, FDR = 0.002) and mast cells (fold difference = 6.6, FDR = 0.002) (**Figure 3C, Supplementary Figure S4A**). While the NK and macrophage (including dendritic cells) clusters were subset for further clustering, the T-cell (106 cells: 65 control, 41 RPL), B-cell (105 cells: 19 control, 86 RPL) and mast cell (26 cells: 4 control, 22 RPL) populations did not possess enough cells for clustering into further phenotypes.

**Figure 3:**
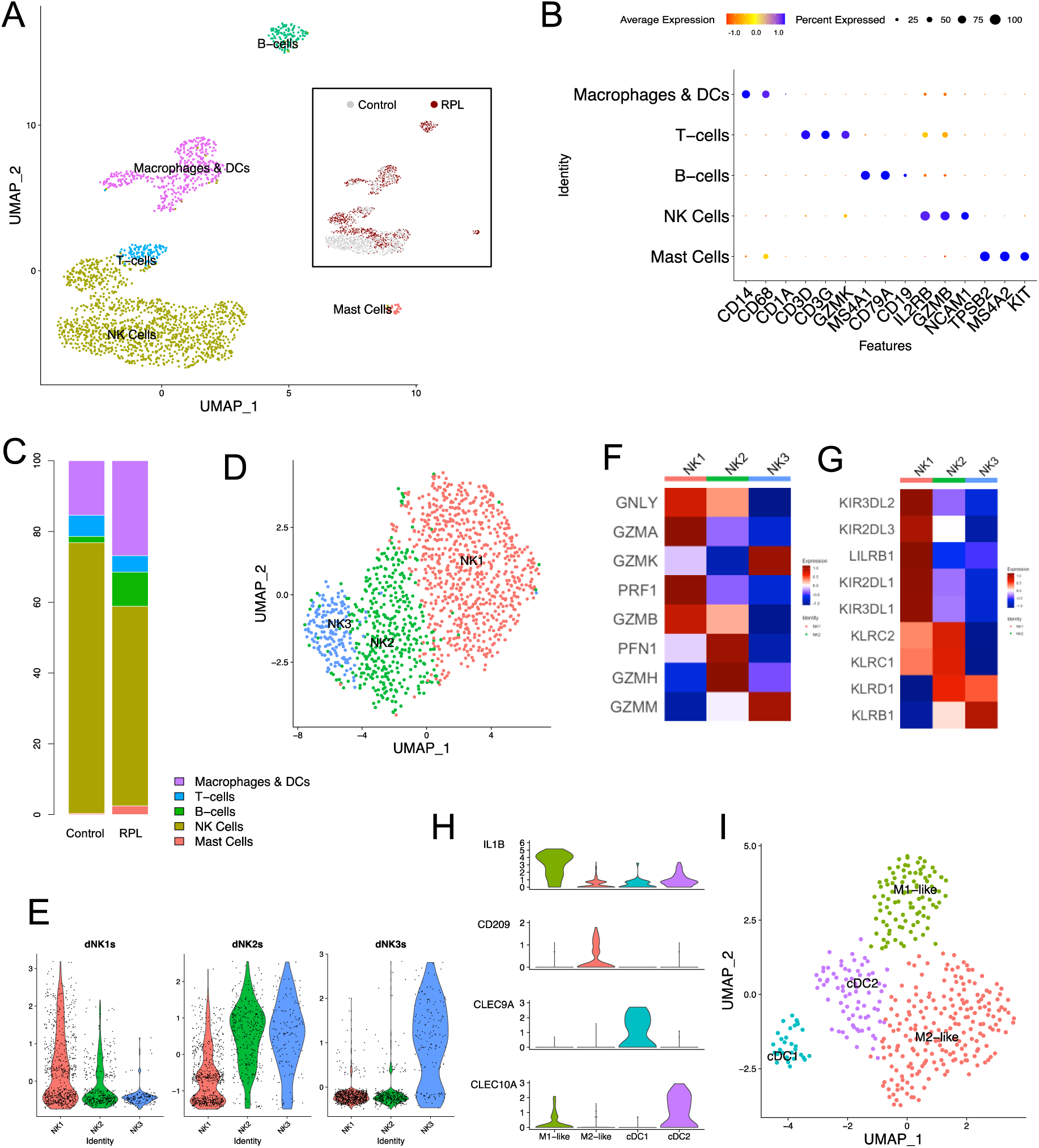
Overview of immune cell populations in the endometrium during the secretory phase. **A)** UMAP of 1,974 immune cells clustered by major immune cell type with inset UMAP showing breakdown of cells by patient groups. **B)** Cell types were identified using known marker gene expression as visualised in dotplot. **C)** Stacked bar chart showing relative proportions of immune cells in both patient groups. **D)** UMAP of subset natural killer cells clustered into three subtypes and identified as previously identified natural killer cells using gene signatures and visualised in violin plots **(E)**. Heatmaps showing the expression of natural killer cell associated genes involved in **F)** cytotoxicity and **G)** immune response. **H)** Violin plots of macrophage and dendritic cell marker gene expression used to identify the sub populations. **I)**. 302 cells formed two macrophage clusters (M1-like and M2-like) and 104 cells formed two conventional dendritic cells (cDC) clusters (cDC1 and cDC2).

### Natural killer cell phenotypes are differentially abundant in control and RPL endometrium

uNK cells are important for immune tolerance at the maternal-fetal interphase, spiral artery remodelling and trophoblast invasion and were the largest cluster of immune cells (identified by *IL2RB*, *GZMB* and *NCAM1* [41]. After further QC, 1,282 cells (795 control, 487 RPL) clustered into 3 subtypes that resembled previously identified decidual NK subtypes [9] (**Figure 3D**) and were named NK1, NK2 and NK3 after the corresponding decidual NK cells (**Figure 3E**). NK1 was the most abundant of the NK cells with 706 cells (464 control, 242 RPL), while the number of cells in NK2 and NK3 sub-clusters was 350 (237 control, 113 RPL) and 148 (45 control, 103 RPL) respectively. NK1 and NK2 were significantly enriched in control patients (p = 0.001 and p = 0.01 respectively, two proportion z-test), while NK3 were significantly enriched in RPL patients (p < 2.2 x 10^-16^, two proportion z-test) (**Figure 1E**). This in part matched observations by Chen *et al* (2021) [42] where NK1 equivalent (cluster 2 dNK) were more abundant in control, and NK3 (cluster 4 dNK) more abundant in RPL.

### Natural killer cells orchestrate cellular changes in the endometrial microenvironment

The expression of granule protein genes (*GZMA*, *GZMB*, *GNLY*, and *PRF1*) were higher in NK1 (**Figure 3F, Supplementary Table S9**) suggesting a more cytotoxic role in the secretory phase. Pathway analysis supports this, with neutrophil degranulation a top GO pathway hit for NK1 but not present in the top 20 most significant hits for NK2 and NK3 (**Supplementary Figure S4B-D**). High levels of KIR receptor genes in NK1 cells (**Figure 3G**) may facilitates communication with the trophoblast if pregnancy occurs [9]. The NK1 cell population likely represents the cytotoxic NK cell type that is responsible for clearance of the SC5 pre-decidual cell type via granule exocytosis [19] (**Supplementary Figure S4B**).

NK2 cells, with significantly upregulated *AREG* and *TNFRSF* genes (**Supplementary Table S9**), resemble a NK population that promotes differentiation of *IGF1*^+^ EnSCs (our SC5) and a *CSF-1*^+^ NK cell population (our NK1) via *AREG* [21]. NK2 cells were in lower proportion in RPL samples (**Figure 1E**), this could explain increased abundance of SC5, and decreased abundance of NK1 in RPL samples.

The NK3 subpopulation expressed significantly higher chemokine ligands and receptor genes including *CCL4L2*, *CCL3*, and *CXCR4* and had upregulated cytokine related and regulation of the immune response pathways (**Supplementary Figure S4D, Supplementary Table S9**). The higher abundance of this cell type in RPL endometrium may explain the increase in immune cells observed in RPL (i.e. NK3 recruits immune cells). All three subtypes of NK cell perform vital roles in the endometrium to facilitate successful implantation and pregnancy (**Supplementary Figure S4E**).

### Macrophage cell phenotypes favour complement activation and the inflammatory response

A cluster of 406 immune cells identified as macrophages and dendritic cells based on marker genes *CD14*, *CD68* and *CD1A* (**Figure 3B**). Four subtypes were identified (**Figure 3H**), two of which were macrophage populations M1-like and M2-like based on known markers of M1 and M2 macrophages, and two were conventional dendritic cells (cDC) cDC1 and cDC2 (**Figure 3I**) that were previously identified as either *CD1C*^+^ or *CD141*^+^ (THBD) [43] (**Supplementary Figure S5A, Supplementary Table S10**). Compared to the M2-like cluster, the M1-like subtype was high in expression for M1 macrophage markers *IL1B* and *CD11c* (*ITGAX*) expression, but low in M2 macrophage marker genes *CD206* (*MRC1*) and *CD209* (*DC-SIGN*) (**Supplementary Figure S5B, Supplementary Table S10**). In pathway analysis using M1-like upregulated genes, the top two pathways from the MsigDB hallmark gene sets were TNFa signalling via NFKB, and inflammatory response, both characteristic of M1 macrophages (**Supplementary** Figure 5C). The M2-like macrophage resembles a *TREM2* high macrophage population identified in melanoma [44]. M2-like macrophages exhibited higher levels of *TREM2* expression compared to M1-like, and of the 14 genes in a gene signature associated with *TREM2* high macrophages, 8 were upregulated compared to M1-like (**Supplementary Figure S5B**). *TREM2* high macrophages are associated with complement activation. In pathway analysis of M2-like cluster upregulated genes, complement activation was the top hit (**Supplementary Figure S5D**). Complement activation is vital at various stages of pregnancy including implantation, and fetal development [45]. Both macrophage populations are likely to be involved in inflammatory signalling, so the higher abundance of both in RPL samples could negatively affect the outcome of pregnancy as high levels of inflammation have been linked to RPL [46].

### Identification of subpopulations of endometrial epithelial and endothelial cells

The epithelial cell population of 717 cells clustered into three subtypes: ciliated epithelial cells (CEC), unciliated epithelial cells 1 (UEC1), and unciliated epithelial cells 2 (UEC2; **Figure 4A**) based on known epithelial marker genes [26]. The CEC cluster (28 cells; 18 control and 10 RPL) was characterised by *HES6*, *FOXJ1*, and *PIFO* gene expression (**Figure 4B**). CECs have been implicated in subfertility with defects in structure and function, and in RPL with altered gene splicing in genes associated with glandular secretions and cilia [47] [48]. The UEC1 (347 cells; 217 control, 130 RPL) and UEC2 (342 cells; 207 control, 135 RPL) clusters, were clearly unciliated based on ciliated cell-associated gene expression, but it was unclear whether either population was glandular or luminal. Both clusters expressed glandular-associated genes *PAEP*, *CXCL14*, and *GPX3*, but luminal cell-associated gene expression was low in both cell types (**Figure 4B, Supplementary Table S11**). Other genes associated with glandular epithelium, *MUC16* and *PAX8* (**Figure 4C,D, Supplementary Table S11**) were upregulated in UEC2 suggesting that they may be glandular epithelium cells. It is possible that sampling and processing of the tissue caused cell death, as although mitochondrial gene content was not particularly high in the cells, the number of reads and genes expressed in UEC1 and UEC2 were markedly lower than in other cell types (**Supplementary Figure S6).**

**Figure 4:**
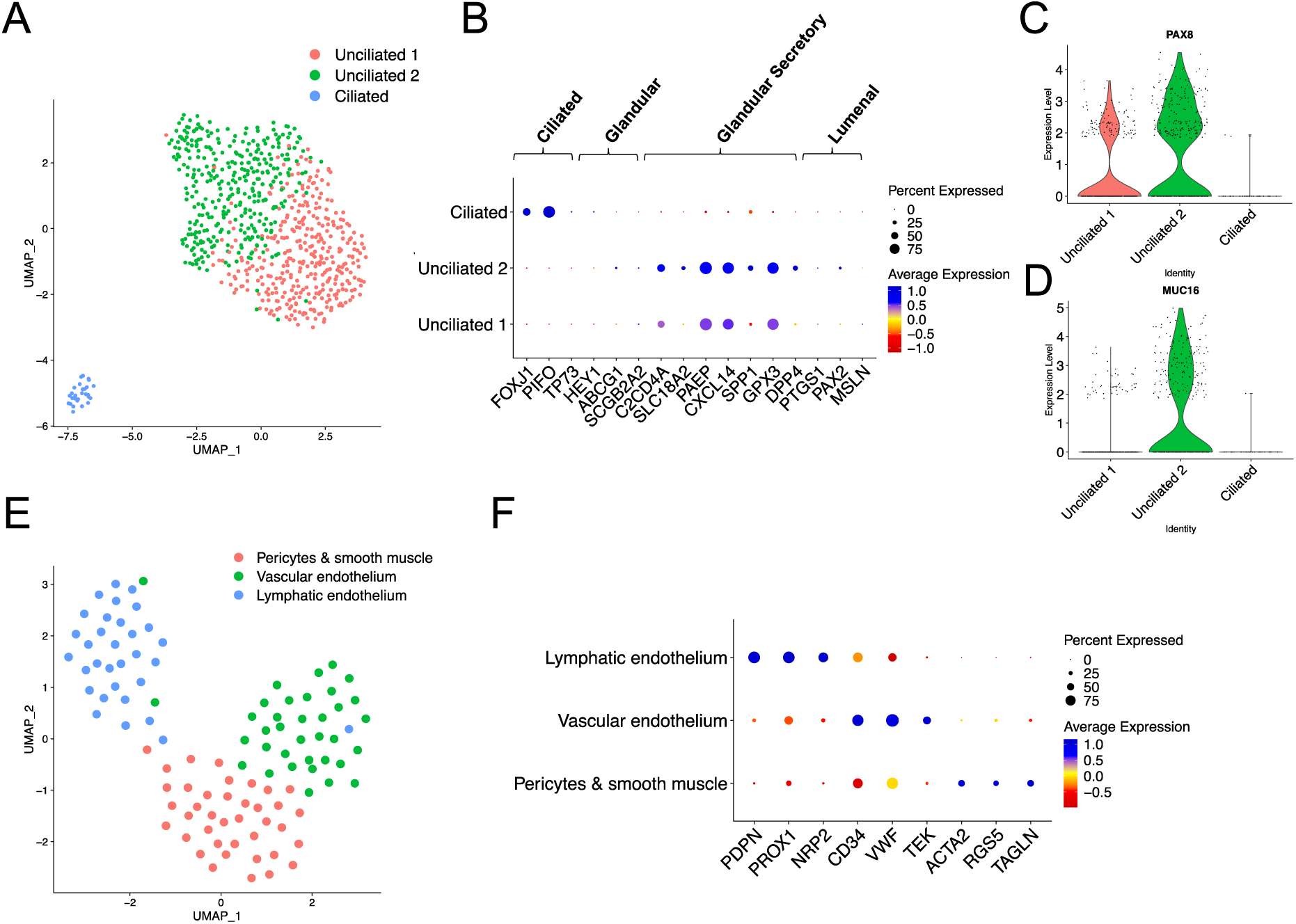
Identification of epithelial and endothelial cell subtypes. **A)** UMAP of 717 epithelial cells designated as ciliated or unciliated using known marker genes of endometrial epithelial cells as visualised in dotplot **(B)**. Violin plots for expression of glandular epithelial marker genes (**C)** PAX8, and (**D)** MUC16. **E)** UMAP of endothelial cells clustered into three subtypes and identified based on the gene expression of known endothelial cell markers **(F)**.

The 107 endothelial cells clustered into three endothelial cell clusters (**Figure 5E**), 39 (17 control, 22 RPL) were pericytes (PC) as identified by of *ACTA2* and *RSG5* gene expression and 36 (20 control, 16 RPL) were vascular endothelium (VE) as identified by *VWF* and *CD34* expression. Lymphatic vessels have been demonstrated in the functionalis and basalis regions of the endometrium [49] and this study observed 32 (14 control, 18 RPL) lymphatic like endothelium (LE) with *PROX1* and *PDPN* expression (**Figure 5F, Supplementary Table S12**).

**Figure 5:**
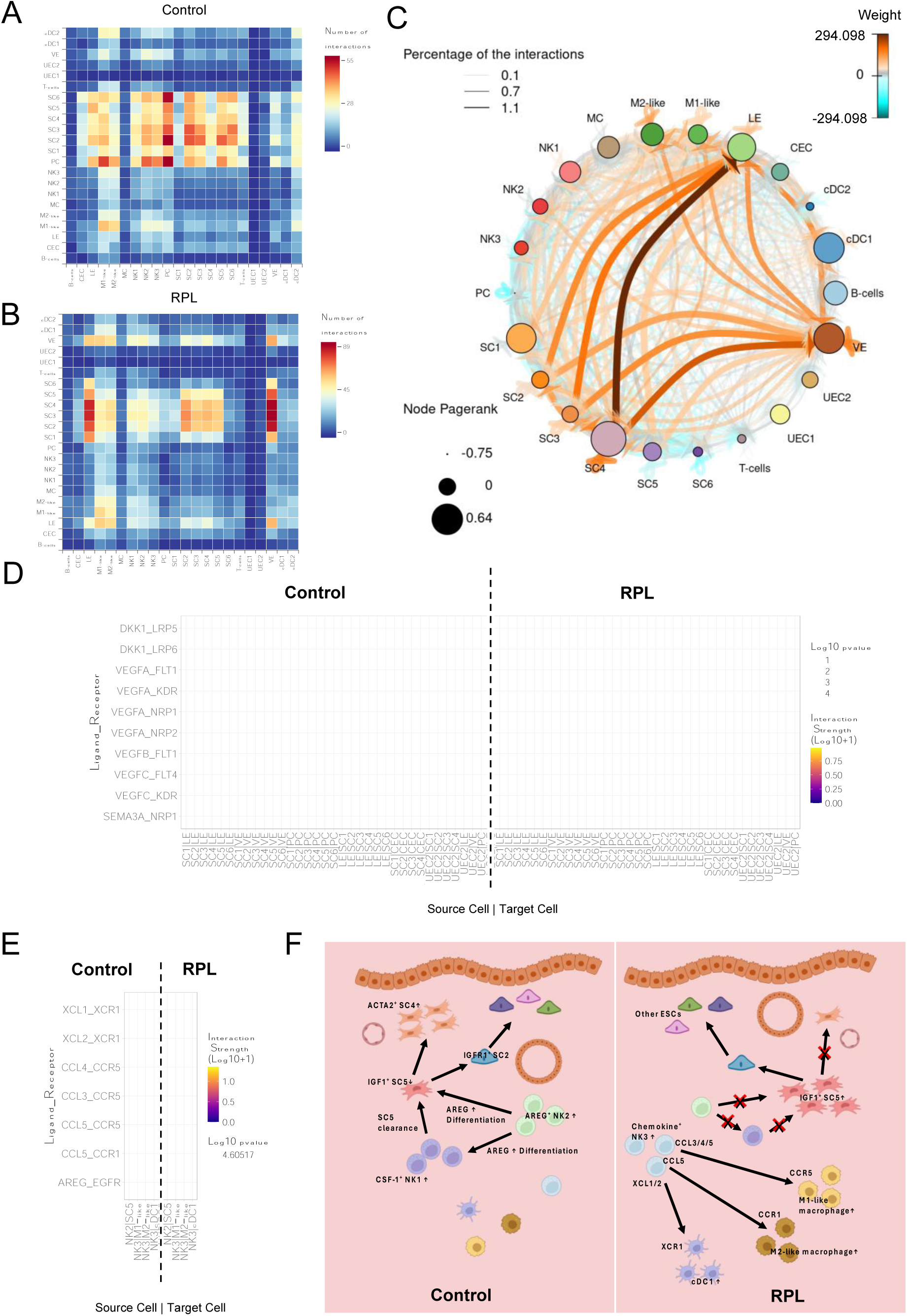
Cell communication reveals differences in processes vital for successful pregnancy. Heatmaps showing the number of interactions between cell types in (**A)** control and (**B)** RPL groups. **C)** Circos plot showing relative interactions between cell types. Dark orange-brown represents higher number of interactions in RPL group, cyan/blue represents higher number of interactions in control group. Width of lines represents total percentage of interactions, and the larger the circle representing cell type the more interactions there are for RPL compared to control (smaller circles represent the opposite – more interactions in control). Dotplot of chosen interactions in Wnt signalling and angiogenesis pathways **(D)** and NK2 and NK3 related interactions of interest **(E)** for both control and RPL groups. The larger the dot, the more significant the interaction, and the more yellow the dot the greater the interaction strength. **F)** Proposed model for differences between control and RPL based on cell type proportion and cell communication differences.

### Cell-cell communication is dysregulated in RPL endometrium

Cell-to-cell communication via ligand and receptor interactions was inferred using CellphoneDB. Despite lower cell numbers, RPL endometrium had more significant cell-cell interactions compared to control (8,426 vs 6,344), consistent with observations in control vs recurrent spontaneous abortion patients [50]. Overall, the greatest cell-cell interaction was between EnSCs and endothelial cells, with increased PC and EnSCs interaction in control endometrium (**Figure 5A**) and increased interactions between LE and VE cells with EnSCs in RPL (**Figure 5B**). B-cells, mast cells, UEC1 and UEC2 were involved in the least number of interactions in both groups. Due to the large number of interactions with differential interaction scores (**Figure 5C**), we investigated those that were significantly different between RPL and control or were only present in one group. Ligand receptor interactions involving genes expressed in biological processes vital for endometrial homeostasis (e.g. Wnt-signalling pathway, angiogenesis, and immune response), and those interactions previously observed in the endometrium were preferentially analysed further. Associations between dysregulation of the Wnt-signalling pathway and angiogenesis, and RPL are well documented [51–54].

### Inappropriate Wnt-signalling may underly unsuccessful pregnancy

There is evidence of canonical (beta-catenin-dependent) and noncanonical (beta-catenin-independent) Wnt-signalling in the endometrium at the mid-secretory phase and during decidualisation. Canonical Wnt-signalling, which regulates cell proliferation, differentiation and maturation [55], is required during EnSC proliferation. However as the mid-secretory phase is reached *DKK1* is upregulated and inhibits canonical Wnt-signalling [56]. This is complicated by the increase in expression of Wnt ligand *WNT4* in EnSCs in response to *BMP-2* and *FOXO1* upregulation [57]. Both *DKK1* and *WNT4* genes were expressed throughout all EnSC subtypes in our dataset, *DKK1* significantly higher in RPL patients compared to control (**Supplementary** Figure 7A). DKK1 repression of canonical Wnt-signalling is apparent in EnSCs as DKK1 acts as a ligand for Wnt co-receptors LRP5 and LRP6, more so in RPL EnSCs as this ligand-receptor interaction is only observed in SC2 and SC6 EnSCs in control patients but in all EnSC subtypes but SC6 in RPL patients (**Supplementary Figure S7B**). In EnSCs, there was also evidence of non-canonical Wnt-signalling, which is involved in cellular polarisation and migration [55], with Wnt ligand WNT5A interacting with receptor ROR2 in SC3 and SC6 in control and SC2, SC3, and SC4 in RPL patients (**Supplementary Figure S7B**). Wnt-signalling between EnSCs and endothelial cells has been previously observed [50], while none was apparent between EnSCs and endothelial cells in our cohort, DKK1-mediated regulation was observed via LRP5/6 almost exclusively in RPL patients (**Figure 5D**) between EnSCs and endothelial cells, and EnSCs and CECs. Wnt-signalling is necessary for successful implantation [58], and given its importance during EnSC proliferation, differentiation, and migration, it would suggest that regulation of the pathway is vital for a successful pregnancy. Differences in the regulation of Wnt-signalling between control and RPL patients, particularly in DKK1-mediated downregulation, as seen in our data, may contribute to unsuccessful pregnancy.

### Dysregulation of Angiogenesis may occur in RPL

Angiogenesis contributes to endometrial remodelling throughout the menstrual cycle and is necessary for successful pregnancy. VEGF associated angiogenesis was investigated by looking at predicted cellular interactions involving VEGF genes (*VEGFA*, *VEGFB*, and *VEGFC)*. VEGFA expressed by EnSCs interacted with FLT1, KDR, NRP1, and NRP2 receptors on endothelial cells, while EnSC VEGFB interacted with endothelial FLT1 receptors [**Figure 5D**]. *VEGFA* regulates angiogenesis while *VEGFB* prevents excessive angiogenesis via inhibition of the *FGF2*/*FGFR1* pathway [59]. *VEGFA* and *VEGFB* facilitated cell communication between EnSCs and endothelial cells was observed in both groups. In RPL samples all EnSC populations were shown to communicate via *VEGFA* and *VEGFB*, while in controls this was only observed in SC2, 5 and 6. VEGFA signalling between UEC2 and EnSCs (SC1-4), and between UEC2 and endothelial cells, was exclusive to RPL samples. Differences in VEGFA/B signalling between control and RPL endometrium observed in this analysis suggest that there is dysregulation of angiogenesis in RPL samples.

Cells with markers characteristic of LE (e.g. *PDPN*) were observed constant with previous studies present in the endometrium but are thought to disappear following the initiation of pregnancy [60]. Regulation of LE may be important for maintaining successful pregnancy. VEGFC and its receptors KDR and especially FLT4 regulate angiogenesis [59]. VEGFC interactions with FLT/KDR were only present in RPL samples between EnSCs and endothelial cells, suggesting potential upregulation of lymphangiogenesis like pathways that could have a detrimental effect on implantation/pregnancy.

### NK cells may be involved in differential abundance of cell types in RPL

A subset of immune cell types was more abundant in RPL samples, potentially recruited by NK3, and was also more abundant in the RPL group. Predicted communication between the NK3 cell and immune cells via appropriate ligand-receptor pairings was often higher or exclusive in RPL cases (**Figure 5E**). Interactions between inflammatory chemokines *CCL3*, *4*, and *5* expressed on NK3 and receptor *CCR5* on M1-like macrophages are exclusively seen in RPL samples, while interactions between *CCL5* from NK3 and receptor *CCR1* on M2-like macrophages was also observed in our data (**Figure 5E**). Interactions between *XCL1* and *2* expressed by NK3 with receptor *XCR1* from cDC1 were also exclusively seen in RPL samples (**Figure 5E**). These interactions would suggest that NK3 recruits other immune cells and the higher abundance of NK3 in RPL samples may explain the increased proportion of immune cells in RPL. The higher abundance of NK3 that express these inflammatory chemokines may also suggest that there are higher levels of inflammation in RPL samples, a condition previously linked to RPL [61].

An interesting interaction was observed between *AREG* expressed by NK2 and receptor *EGFR* by SC5. *AREG* has been associated with differentiation of fibroblasts to myofibroblasts via *EGFR* [62, 63]. While NK cells have traditionally been associated with clearance of fibroblasts [64], in the endometrium NK subtypes may have a role in the differentiation of pre-decidualised EnSC (SC5) to the myofibroblast phenotype (SC4) via this *AREG*-*EGFR* axis. This would be consistent with the differential abundances of NK2, SC4, and SC5 in control samples.

### NK cells orchestrate differences in cell type abundance in control and RPL

Cell communication analysis of our data suggests that both the Wnt-signalling pathway and angiogenesis may be dysregulated in our RPL cohort, which has been previously been observed [51–54]. Perhaps more interesting and less explored, was the difference in cellular composition in control and RPL endometrium microenvironments.

During the window of implantation, the endometrium is populated with the multi-functional mesenchymal/fibroblast-like EnSCs. Our data suggest that these EnSCs undergo differentiation into sub-populations of “decidualised” EnSCs that, while displaying similar gene expression profiles, potentially perform different roles in the endometrium. Of the six subpopulations SC5, significantly more abundant in RPL patients, appears to be the EnSC that initiates decidualisation and may differentiate into a myofibroblast-like EnSC SC4, which is significantly more abundant in control patients. Differences in abundance could be explained by the increased number of NK1 and NK2 cells in control patients. NK2 potentially has roles in promoting differentiation of EnSCs, via *AREG* expression, to myofibroblasts, and NKs to the NK1 subtype, a subtype of NK cell involved with the clearance of EnSCs via degranulation. The NK3 subtype was more abundant in RPL patients and displayed marker genes that are involved in the recruitment of immune cells, which may explain the significantly greater abundance of immune cells in RPL samples.

Along with potential dysregulation of angiogenesis and the Wnt-signalling pathway, two processes vital to a successful pregnancy, the different cell composition between the two patient groups provides a model (**Figure 5F**) whereby NK cell populations orchestrate environments conducive, or otherwise, to a successful pregnancy.

## Conclusion

This study provides compelling evidence of cellular differences in the endometrium between women who have experienced RPL and those who have not, offering potential insights into endometrial mechanisms that may contribute to RPL. Despite the clear evidence that there are multiple EnSC subtypes, consistent with previous literature, there are limitations associated with these analyses including sample size and patient heterogeneity. In addition, while we found supporting evidence of all our EnSC populations in the broader literature, no one study contained all SC1-6. And there was little agreement with regard to EnSC phenotypes. This may reflect varying methods for cell dissociation, which can differentially affect cell transcriptomes [65]. Differences in the menstrual cycle phase may also contribute to these differences [66, 67]. Future studies could investigate whether cell differentiation pathways are replicated in culture which would allow mechanisms to be studies in more detail and specific potential interventions to be tested.

In summary, this study provides valuable new insights into endometrial cellular differences in RPL, paving the way for future research to further explore these mechanisms. With larger, more consistent samples, we can deepen our understanding of EnSC subtypes and move closer to identifying potential therapeutic interventions.

## Methods

### Ethics

The study was approved by Isle of Wight, Portsmouth & South East Hampshire Research Ethics Committee (08/H0502/162). Written informed consent was given by all participants.

### Endometrial biopsy collection

Participants that met the study criteria were recruited for collection of an endometrial tissue biopsy at a tertiary fertility and gynaecology referral centre in Southampton, UK. These included recurrent pregnancy loss and control participants. Control participants (n = 4) were recruited from healthy fertile women who elected to donate eggs at the local fertility centre in Southampton having met the criteria for egg donation. Control participants had no history of subfertility or recurrent pregnancy loss. Recurrent pregnancy loss participants (n = 3) had a history of three or more first trimester losses [1].

All endometrial biopsies were collected from women experiencing normal menstrual cycles without hormonal stimulation aged between 29 and 43 years and not suffering from premature ovarian failure or any infective processes. Endometrial biopsies were collected on days 21-24 of the menstrual cycle: 7-10 days post the luteinising hormone surge, encompassing the window of implantation (LH + 7 - 10). Endometrial biopsies (approximately 1.5 cm x 0.5 cm) were collected by a Pipelle sampler (Laboratorie C.C.D) [68].

### Single cell suspension

Endometrial biopsies were collected into culture media (50:50 DMEM/ Ham’s F12 nutrient mixture, containing 5% streptomycin, no FBS) at room temperature, and tissue digestion was started within 1 h of tissue collection. The biopsy was minced into 1-2 mm^3^ pieces and digested with 0.7 mg/ml type 1A collagenase (Sigma C9891) in culture media at 37°C for 45 min with gentle agitation every 15 min. Following 5 min at room temperature the digested endometrial cell suspension was filtered three times through a 30 µm sieve and rinsed with culture media. The cell suspension was centrifuged at 1000 rpm for 10 min and resuspended in 2 mL 0.04 mg/ml BSA in PBS. Cells were counted with trypan blue using a hemocytometer, aiming for 1000 cells/ml.

### Droplet-based scRNA-seq

Samples were processed using the 10 x Genomics Chromium Next GEM Single-Cell 3ʹ GEM, Library & Gel Bead Kit v3.1 and Chromium Controller instrument (10 x Genomics) following the manufacturer’s protocol. Cell suspensions of 5000–10,000 single cells were loaded to each channel of a chromium single-cell chip G along with reverse transcription master mix and single-cell 3-gel beads, with the average recovery rate of 2000 cells. The samples were processed using 10 × Genomics v3.1 barcoding chemistry kits. Following cell lysis, first-strand cDNA synthesis and amplification were carried out according to the manufacturer’s instructions. A Single-cell 3’-Library was prepared and 150bp PE sequencing carried out using an Illumina novaseq6000 (Oxford Genomics Centre).

### Single-cell gene expression quantification

The Cell Ranger software pipeline (v3.1.0) provided by 10 × Genomics was used to demultiplex cellular barcodes, map reads to the genome and transcriptome using the STAR aligner, and down-sample reads as required to generate normalised aggregate data across samples, producing a matrix of gene counts versus cell. Seurat (v4.1.0) was used to generate Seurat objects for each sample.

### Filtering and preprocessing

For each sample, cells with fewer than 500 genes were considered to be of low-quality and removed. R package DoubletFinder (V2.0) [69] was used to remove cells that were considered doublets. Cells were also removed if the number of genes detected was greater than 6,000, percentage of mitochondrial gene content was greater than 25%, or log10GenesPerUmi was greater than 0.8. Seurat (v4.1.0) [70] was used to integrate the data for each patient after this initial quality control and the final data set consisted of 10,022 cells from seven samples (sample GX47 was removed due to poor quality of data). Epithelial, immune, stromal, and endothelial cell populations were identified using cell-specific markers and *a priori* knowledge, and subset into individual Seurat objects for further analysis.

### Cell-type identification

After clustering of the 2,066 immune cells, 92 cells were considered contaminated/doublets and the remaining cells were further subset based on known marker genes into natural killer cells (NK), macrophages and dendritic cells, T-cells, B-cells, and mast cells. All 8 cell types underwent further clustering after SCTransform was used on each object to regress out differences caused by mitochondrial gene content, the total number of UMIs, and by individual patient:

#### Endometrial stromal cells (EnSC)

7,109 EnSCs were clustered using the top 30 principal components, k.parameter of 50, and resolution of 0.5. Two clusters displayed high expression of marker genes associated with natural killer cells, and haemoglobin genes so were removed from the analysis resulting in a total of 6,501 cells for further analysis. The Seurat object was subset into RPL and control EnSCs and clustered separately. Again SCTransform was used to regress out differences caused by mitochondrial gene content, the total number of UMIs, and by individual patient for both subsets. Clustering was performed for the RPL and control EnSCs using the top 8 principal components, a k parameter of 60, and a resolution of 0.5. After naming the clusters for the individual subsets, the RPL and control objects were merged and reclustered. SCTransform was used to regress out differences caused by mitochondrial gene content, the total number of UMIs, and by individual patient. The top 20 principal components, a k.parameter of 50, and a resolution of 0.5 were used to cluster the EnSCs into 6 individual clusters. Seurat function “FindAllMarkers” was used to identify genes that were upregulated in each EnSC cluster that were used to determine possible functions and compare with previously identified EnSC subtypes.

#### Natural killer cells

1,331 NK cells were clustered using the top 12 principal components (PCs), a k parameter of 10, and a resolution of 0.5. A subset of cells were removed as suspected doublets due to expression of macrophage gene markers leaving 1,282 cells for further analysis. Cells were reclustered (top 12 PCs, k parameter of 60, resolution of 0.4) giving 3 clusters which were identified using gene marker lists for 3 previously identified NK cell types [9]. Each cell was scored using Seurat function “AddModuleScore” and visualised by cluster using “VlnPlot”.

#### Macrophages and dendritic cells

406 cells designated macrophage and dendritic cells. Cells were clustered using the top 12 principal components, a k parameter of 0.6, and a resolution of 1.5 which gave 7 clusters. Following marker gene identification using “FindAllMarkers” it was determined that the cells had been over clustered and reduced to 4 by combining similar clusters (i.e. those expressing the same marker genes).

#### B-cells, T-cells, and mast cells

237 immune cells clustered into B-cells (105 cells), T-cells (106), and mast cells (26). Each subset was too small to stratify into different phenotypes of each cell and were not clustered any further.

#### Epithelial cells

740 epithelial cells were clustered using the top 12 principal components, a k parameter of 30, and resolution of 0.8. After running “FindAllMarkers” a small cluster was determined to be contaminated with EnSCs and removed leaving 717 cells which were reclustered using the top 12 principal components, k parameter of 50, and resolution of 0.6. FindAllMarkers was run again, and the genes most upregulated in each cluster were used to determine which type of epithelial cell each was.

#### Endothelial cells

107 endothelial cells were clustered using the first 9 principal components, k parameter of 30, and resolution of 1. Upregulated genes in each cluster were used to identify individual endothelial cell types.

### Data analysis

#### Cell proportion differences

A comparison of the proportion of cells present in control and RPL was made using R package scProportionTest. Number of bootstraps used in permutation_test was set to 10,000.

#### Pathway analysis

Pathway analysis was performed on genes that were considered upregulated for a cell type based on genes identified by Seurat function “FindAllMarkers” using R package enrichR (v3.1) considering gene ontology biological processes and MsigDB hallmark gene sets.

#### Trajectory analysis

For the EnSC subset Monocle3 was used to infer cell trajectories based on the IGF1+ SC5 population being the root node (based on literature).

#### Transcription factor activity

Raw count matrices and pySCENIC were used to infer transcription factor activity in all cells. Motif collection mc9nr was used. Wilcoxon rank sum test was used to determine significant differentially activated transcription factors in EnSC populations based on a log2 fold change > 0.1 and an adjusted p value < 0.05. The binarization.py script from pySCENIC (v0.12.1) was used to identify cells that were considered “ON” and “OFF” for each transcription factor.

#### Validation data sets

Single cell RNA-seq data for the endometrium was obtained from Wang *et al* (2020) [4] and Garcia-Alonso *et al* (2021) [26]. Cells identified as EnSCs and that were from the mid secretory phase were subset from each data set. Both sets of EnSC were integrated using canonical correlation analysis (CCA), and clustered using the top 30 principal components, k parameter of 10, and a resolution of 0.6. “FindAllMarkers” was used to identify genes upregulated in each cluster which were used for comparison with EnSC clusters from our data set.

#### Cell communication

CellphoneDB (v4.1.0) was used to identify potential interaction across cell types in the control and RPL endometrium. A threshold of 25% was used as a minimum number of cells of an individual cell type that a ligand or receptor had to be expressed in to be included in analysis. A p value of < 0.05 based on 1,000 permutations was used to determine if an interaction was significant. We then selected interactions that were deemed biologically relevant with regards to the endometrium.

## Data availability

Raw sequencing data will be available on GEO upon publication.

## Acknowledgements

We would like to thank the women who consented to be part of this study and clinical staff who assisted with sample collection. The authors acknowledge the use of the IRIDIS High Performance Computing Facility, and associated support services at the University of Southampton, in the completion of this work, and the Faculty of Medicine BIO-R, at the University of Southampton for their support and assistance in this work. Funding from Wellbeing of Women.

## References

1. Obstetricians, R.C.o. and Gynaecologists, Green-top Guideline Number 17. The Investigation and Treatment of Couples with Recurrent First-trimester and Second-trimester Miscarriage. 2011.

2. Rai, R. and L. Regan, Recurrent miscarriage. Lancet, 2006. 368(9535): p. 601–11.

3. Ogasawara, M., et al., Embryonic karyotype of abortuses in relation to the number of previous miscarriages. Fertil Steril, 2000. 73(2): p. 300–4.

4. Wang, W., et al., Single-cell transcriptomic atlas of the human endometrium during the menstrual cycle. Nat Med, 2020. 26(10): p. 1644–1653.

5. Díaz-Gimeno, P., et al., A genomic diagnostic tool for human endometrial receptivity based on the transcriptomic signature. Fertil Steril, 2011. 95(1): p. 50–60, 60.e1-15.

6. Enciso, M., et al., Development of a new comprehensive and reliable endometrial receptivity map (ER Map/ER Grade) based on RT-qPCR gene expression analysis. Hum Reprod, 2018. 33(2): p. 220–228.

7. Díaz-Gimeno, P., et al., The accuracy and reproducibility of the endometrial receptivity array is superior to histology as a diagnostic method for endometrial receptivity. Fertil Steril, 2013. 99(2): p. 508–17.

8. Hernández-Vargas, P., M. Muñoz, and F. Domínguez, Identifying biomarkers for predicting successful embryo implantation: applying single to multi-OMICs to improve reproductive outcomes. Hum Reprod Update, 2020. 26(2): p. 264–301.

9. Vento-Tormo, R., et al., Single-cell reconstruction of the early maternal-fetal interface in humans. Nature, 2018. 563(7731): p. 347–353.

10. Filant, J. and T.E. Spencer, Uterine glands: biological roles in conceptus implantation, uterine receptivity and decidualization. Int J Dev Biol, 2014. 58(2-4): p. 107–16.

11. Olmos-Ortiz, A., et al., Innate Immune Cells and Toll-like Receptor-Dependent Responses at the Maternal-Fetal Interface. Int J Mol Sci, 2019. 20(15).

12. Tabib, T., et al., Myofibroblast transcriptome indicates SFRP2(hi) fibroblast progenitors in systemic sclerosis skin. Nat Commun, 2021. 12(1): p. 4384.

13. Werner, S., et al., MRTF-A controls myofibroblastic differentiation of human multipotent stromal cells and their tumour-supporting function in xenograft models. Sci Rep, 2019. 9(1): p. 11725.

14. Small, E.M., The actin-MRTF-SRF gene regulatory axis and myofibroblast differentiation. J Cardiovasc Transl Res, 2012. 5(6): p. 794–804.

15. Queckbörner, S., et al., Stromal Heterogeneity in the Human Proliferative Endometrium-A Single-Cell RNA Sequencing Study. J Pers Med, 2021. 11(6).

16. Zheng, Y., et al., Characterization of placental and decidual cell development in early pregnancy loss by single-cell RNA sequencing. Cell Biosci, 2022. 12(1): p. 168.

17. Eremichev, R., et al., Scar-Free Healing of Endometrium: Tissue-Specific Program of Stromal Cells and Its Induction by Soluble Factors Produced After Damage. Front Cell Dev Biol, 2021. 9: p. 616893.

18. Qian, J., et al., A pan-cancer blueprint of the heterogeneous tumor microenvironment revealed by single-cell profiling. Cell Res, 2020. 30(9): p. 745–762.

19. Lucas, E.S., et al., Recurrent pregnancy loss is associated with a pro-senescent decidual response during the peri-implantation window. Commun Biol, 2020. 3(1): p. 37.

20. Brighton, P.J., et al., Clearance of senescent decidual cells by uterine natural killer cells in cycling human endometrium. Elife, 2017. 6.

21. Shi, J.W., et al., An IGF1-expressing endometrial stromal cell population is associated with human decidualization. BMC Biol, 2022. 20(1): p. 276.

22. Shih, A.J., et al., Single-cell analysis of menstrual endometrial tissues defines phenotypes associated with endometriosis. BMC Med, 2022. 20(1): p. 315.

23. Franasiak, J.M., et al., Endometrial CXCL13 expression is cycle regulated in humans and aberrantly expressed in humans and Rhesus macaques with endometriosis. Reprod Sci, 2015. 22(4): p. 442–51.

24. Hayashi, K., et al., WNTs in the neonatal mouse uterus: potential regulation of endometrial gland development. Biol Reprod, 2011. 84(2): p. 308–19.

25. Cornillie, F.J., J.M. Lauweryns, and I.A. Brosens, Normal human endometrium. An ultrastructural survey. Gynecol Obstet Invest, 1985. 20(3): p. 113–29.

26. Garcia-Alonso, L., et al., Mapping the temporal and spatial dynamics of the human endometrium in vivo and in vitro. Nat Genet, 2021. 53(12): p. 1698–1711.

27. Mericskay, M., J. Kitajewski, and D. Sassoon, Wnt5a is required for proper epithelial-mesenchymal interactions in the uterus. Development, 2004. 131(9): p. 2061–72.

28. Miller, C., A. Pavlova, and D.A. Sassoon, Differential expression patterns of Wnt genes in the murine female reproductive tract during development and the estrous cycle. Mech Dev, 1998. 76(1-2): p. 91–9.

29. Hayashi, K., et al., Wnt genes in the mouse uterus: potential regulation of implantation. Biol Reprod, 2009. 80(5): p. 989–1000.

30. Tulac, S., et al., Dickkopf-1, an inhibitor of Wnt signaling, is regulated by progesterone in human endometrial stromal cells. J Clin Endocrinol Metab, 2006. 91(4): p. 1453–61.

31. Shen, Y., et al., Epigenome-Wide Association Study Indicates Hypomethylation of MTRNR2L8 in Large-Artery Atherosclerosis Stroke. Stroke, 2019. 50(6): p. 1330–1338.

32. Meng, W., et al., Anti-apoptotic genes and non-coding RNAs are potential outcome predictors for ulcerative colitis. Funct Integr Genomics, 2023. 23(2): p. 165.

33. Kokawa, K., T. Shikone, and R. Nakano, Apoptosis in the human uterine endometrium during the menstrual cycle. J Clin Endocrinol Metab, 1996. 81(11): p. 4144–7.

34. Critchley, H.O., et al., Role of inflammatory mediators in human endometrium during progesterone withdrawal and early pregnancy. J Clin Endocrinol Metab, 1999. 84(1): p. 240–8.

35. Workel, H.H., et al., A Transcriptionally Distinct CXCL13(+)CD103(+)CD8(+) T-cell Population Is Associated with B-cell Recruitment and Neoantigen Load in Human Cancer. Cancer Immunol Res, 2019. 7(5): p. 784–796.

36. Kitaya, K. and T. Yasuo, Aberrant expression of selectin E, CXCL1, and CXCL13 in chronic endometritis. Mod Pathol, 2010. 23(8): p. 1136–46.

37. Murphy, D.J., et al., Distinct thresholds govern Myc’s biological output in vivo. Cancer Cell, 2008. 14(6): p. 447–57.

38. Wan, S., et al., METTL3-dependent m(6)A methylation facilitates uterine receptivity and female fertility via balancing estrogen and progesterone signaling. Cell Death Dis, 2023. 14(6): p. 349.

39. Johnson, M.C., et al., Augmented cell survival in eutopic endometrium from women with endometriosis: expression of c-myc, TGF-beta1 and bax genes. Reprod Biol Endocrinol, 2005. 3: p. 45.

40. Chou, Y.C., et al., Killer cell immunoglobulin-like receptors (KIR) and human leukocyte antigen-C (HLA-C) allorecognition patterns in women with endometriosis. Sci Rep, 2020. 10(1): p. 4897.

41. Smith, S.L., et al., Diversity of peripheral blood human NK cells identified by single-cell RNA sequencing. Blood Adv, 2020. 4(7): p. 1388–1406.

42. Chen, P., et al., The Immune Atlas of Human Deciduas With Unexplained Recurrent Pregnancy Loss. Front Immunol, 2021. 12: p. 689019.

43. MacDonald, K.P., et al., Characterization of human blood dendritic cell subsets. Blood, 2002. 100(13): p. 4512–20.

44. Xiong, D., Y. Wang, and M. You, A gene expression signature of TREM2(hi) macrophages and γδ T cells predicts immunotherapy response. Nat Commun, 2020. 11(1): p. 5084.

45. Girardi, G., et al., Essential Role of Complement in Pregnancy: From Implantation to Parturition and Beyond. Front Immunol, 2020. 11: p. 1681.

46. Wu, Z., et al., Pro-Inflammatory Signature in Decidua of Recurrent Pregnancy Loss Regardless of Embryonic Chromosomal Abnormalities. Front Immunol, 2021. 12: p. 772729.

47. Pearson-Farr, J.E., et al., Ultrastructural cilia defects in multi-ciliated uterine glandular epithelial cells from women with reproductive failure. Reproduction, 2024. 167(1).

48. Pearson-Farr, J., et al., Endometrial gland specific progestagen-associated endometrial protein and cilia gene splicing changes in recurrent pregnancy loss. Reprod Fertil, 2022. 3(3): p. 162–72.

49. Donoghue, J.F., et al., Lymphangiogenesis of normal endometrium and endometrial adenocarcinoma. Hum Reprod, 2007. 22(6): p. 1705–13.

50. Du, L., et al., Single-cell transcriptome analysis reveals defective decidua stromal niche attributes to recurrent spontaneous abortion. Cell Prolif, 2021. 54(11): p. e13125.

51. Chen, C., et al., Endometrial protein expression and phosphorylation landscape decipher aberrant insulin and mTOR signalling in patients with recurrent pregnancy loss. Reprod Biomed Online, 2024. 48(1): p. 103585.

52. Chronopoulou, E., et al., Wnt4, Wnt6 and β-catenin expression in human placental tissue - is there a link with first trimester miscarriage? Results from a pilot study. Reprod Biol Endocrinol, 2022. 20(1): p. 51.

53. Zhang, Q. and J. Yan, Update of Wnt signaling in implantation and decidualization. Reprod Med Biol, 2016. 15(2): p. 95–105.

54. Sultana, S., et al., Are Altered Expression of Vascular Endothelial Growth Factor and Placental Growth Factor Associated with Placental Angiogenesis in Recurrent Pregnancy Loss? Indian J Clin Biochem, 2024. 39(3): p. 387–391.

55. Qin, K., et al., Canonical and noncanonical Wnt signaling: Multilayered mediators, signaling mechanisms and major signaling crosstalk. Genes Dis, 2024. 11(1): p. 103–134.

56. Liu, Y., et al., Excessive ovarian stimulation up-regulates the Wnt-signaling molecule DKK1 in human endometrium and may affect implantation: an in vitro co-culture study. Hum Reprod, 2010. 25(2): p. 479–90.

57. Li, Q., et al., WNT4 acts downstream of BMP2 and functions via β-catenin signaling pathway to regulate human endometrial stromal cell differentiation. Endocrinology, 2013. 154(1): p. 446–57.

58. Sonderegger, S., J. Pollheimer, and M. Knöfler, Wnt signalling in implantation, decidualisation and placental differentiation--review. Placenta, 2010. 31(10): p. 839–47.

59. Shibuya, M., Vascular Endothelial Growth Factor (VEGF) and Its Receptor (VEGFR) Signaling in Angiogenesis: A Crucial Target for Anti- and Pro-Angiogenic Therapies. Genes Cancer, 2011. 2(12): p. 1097–105.

60. Volchek, M., et al., Lymphatics in the human endometrium disappear during decidualization. Hum Reprod, 2010. 25(10): p. 2455–64.

61. Yang, X., et al., The Update Immune-Regulatory Role of Pro- and Anti-Inflammatory Cytokines in Recurrent Pregnancy Losses. Int J Mol Sci, 2022. 24(1).

62. Zaiss, D.M.W., et al., Emerging functions of amphiregulin in orchestrating immunity, inflammation, and tissue repair. Immunity, 2015. 42(2): p. 216–226.

63. Zhao, R., et al., Sustained amphiregulin expression in intermediate alveolar stem cells drives progressive fibrosis. Cell Stem Cell, 2024. 31(9): p. 1344–1358.e6.

64. Fasbender, F., et al., Natural Killer Cells and Liver Fibrosis. Front Immunol, 2016. 7: p. 19.

65. van den Brink, S.C., et al., Single-cell sequencing reveals dissociation-induced gene expression in tissue subpopulations. Nat Methods, 2017. 14(10): p. 935–936.

66. Gellersen, B. and J.J. Brosens, Cyclic decidualization of the human endometrium in reproductive health and failure. Endocr Rev, 2014. 35(6): p. 851–905.

67. Edgell, T.A., L.J. Rombauts, and L.A. Salamonsen, Assessing receptivity in the endometrium: the need for a rapid, non-invasive test. Reprod Biomed Online, 2013. 27(5): p. 486–96.

68. Stocker, L., F. Cagampang, and Y. Cheong, Identifying stably expressed housekeeping genes in the endometrium of fertile women, women with recurrent implantation failure and recurrent miscarriages. Sci Rep, 2017. 7(1): p. 14857.

69. McGinnis, C.S., L.M. Murrow, and Z.J. Gartner, DoubletFinder: Doublet Detection in Single-Cell RNA Sequencing Data Using Artificial Nearest Neighbors. Cell Syst, 2019. 8(4): p. 329–337.e4.

70. Hao, Y., et al., Integrated analysis of multimodal single-cell data. Cell, 2021. 184(13): p. 3573–3587.e29.

