## Supplementary Figure 1 for "Changes in the cellular composition of the endometrium during the implantation window are associated with recurrent pregnancy loss"

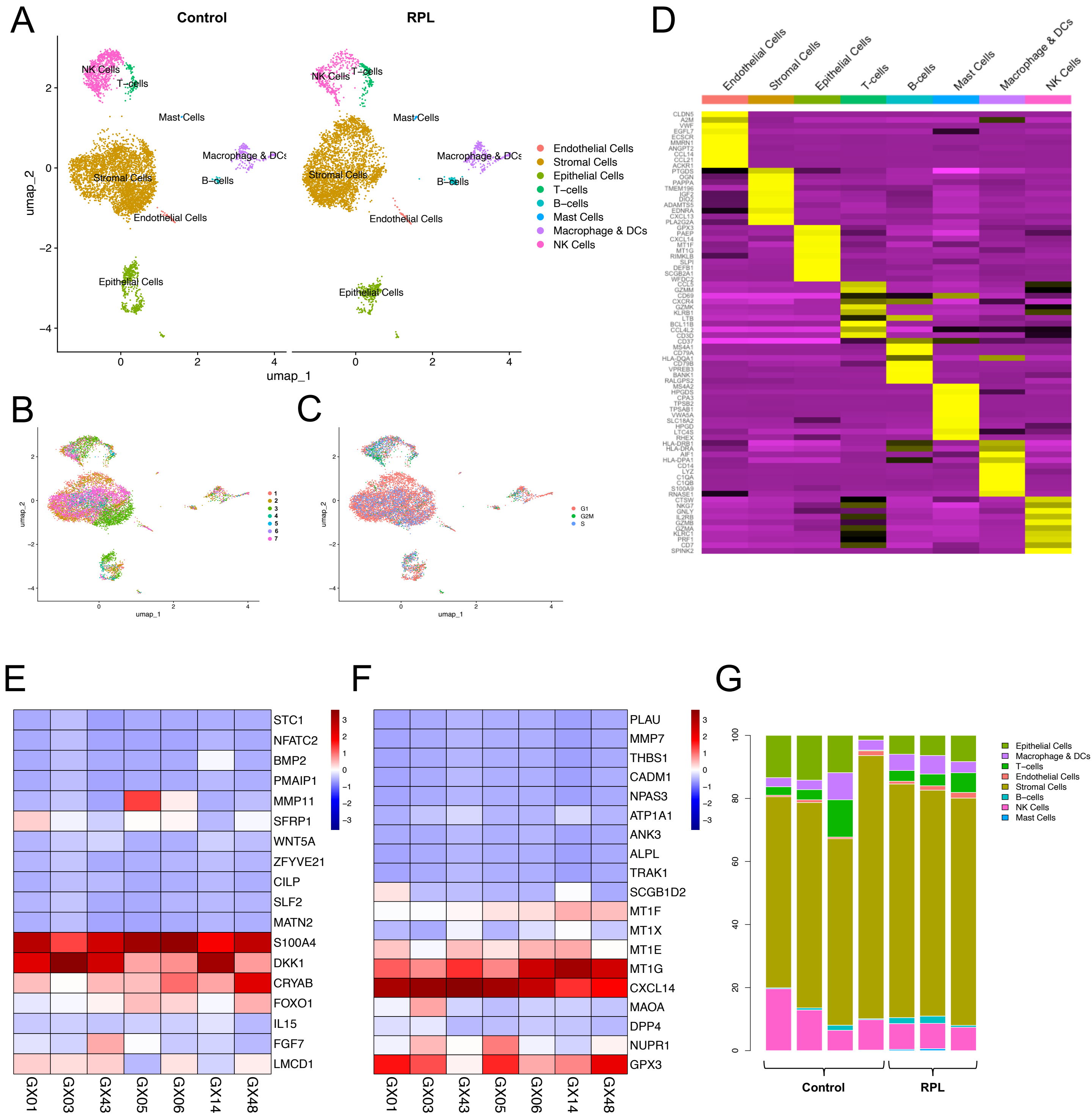


**Supplementary Figure S1 Single cell profiling of control and RPL endometrium.** UMAPs of main cell types for control and RPL groups **(A)**, cell breakdown by patient **(B)**, and cell breakdown by cell cycle phase **(C)**. **D)** Heatmap of top 10 most upregulated genes in each cell type compared to other cell types. Heatmaps showing relative gene expression of genes known to be expressed at various stages of the menstrual cycle in **E)** endometrial stromal cells, and **F)** endometrial epithelial cells. **G)** Major cell proportions by patient.
