## Supplementary Figure 2 for "Changes in the cellular composition of the endometrium during the implantation window are associated with recurrent pregnancy loss"

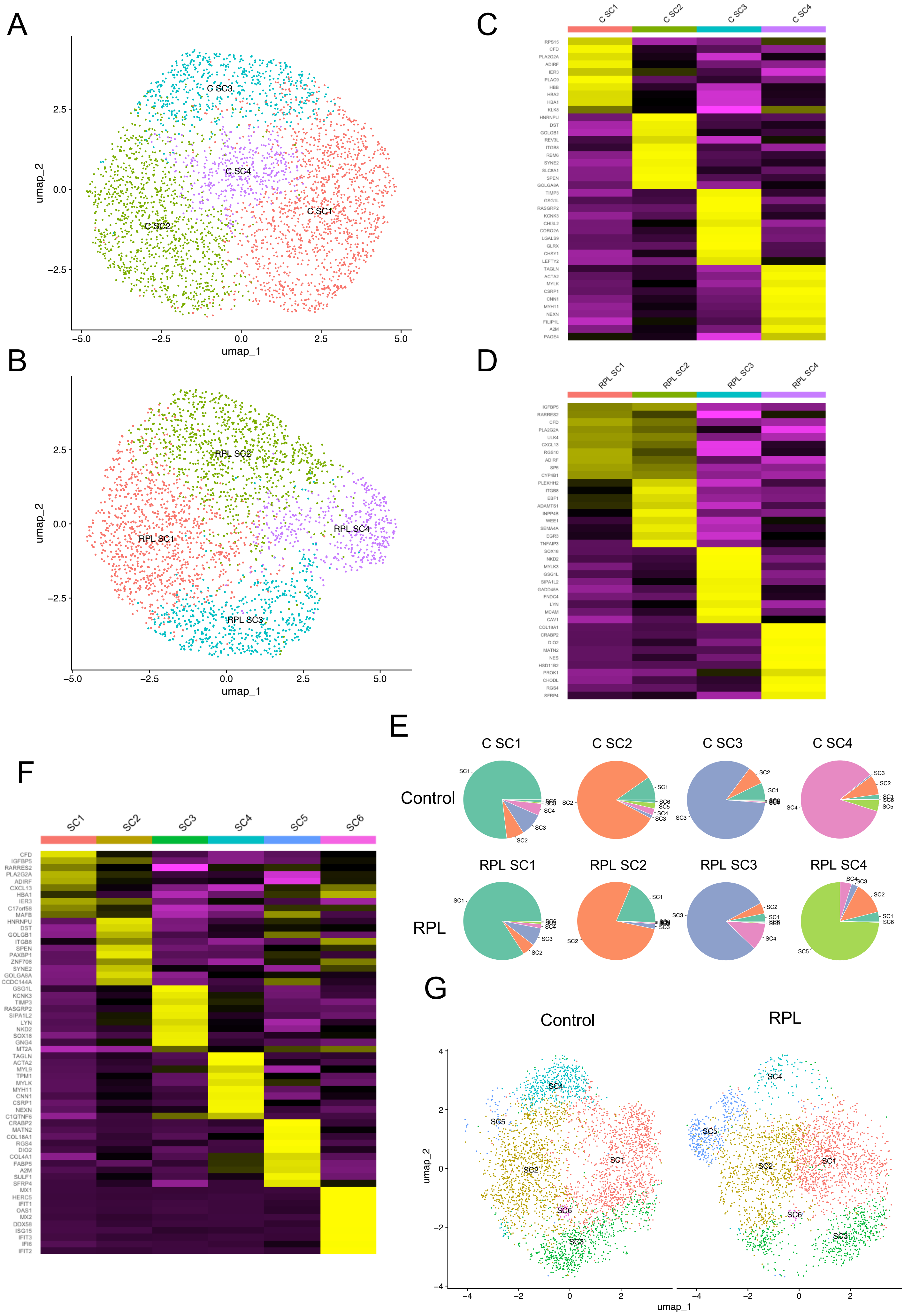


**Supplementary Figure S2 Endometrial stromal cell populations in control and RPL groups.** UMAPs for clustering of endometrial stromal cells (EnSCs) in **A)** control and **B)** RPL patient groups. Heatmaps of the top 10 most upregulated genes in EnSC populations in **C)** control and **D)** RPL patient groups. **E)** Pie charts showing how EnSCs by cluster in both groups re-cluster proportionally when both groups are combined for analysis. **F)** Heatmap showing top 10 most upregulated genes in all EnSC subtypes in the cobined data set. **G)** UMAPs of clustered EnSCs for both groups.
