## Supplementary Figure 3 for "Changes in the cellular composition of the endometrium during the implantation window are associated with recurrent pregnancy loss"

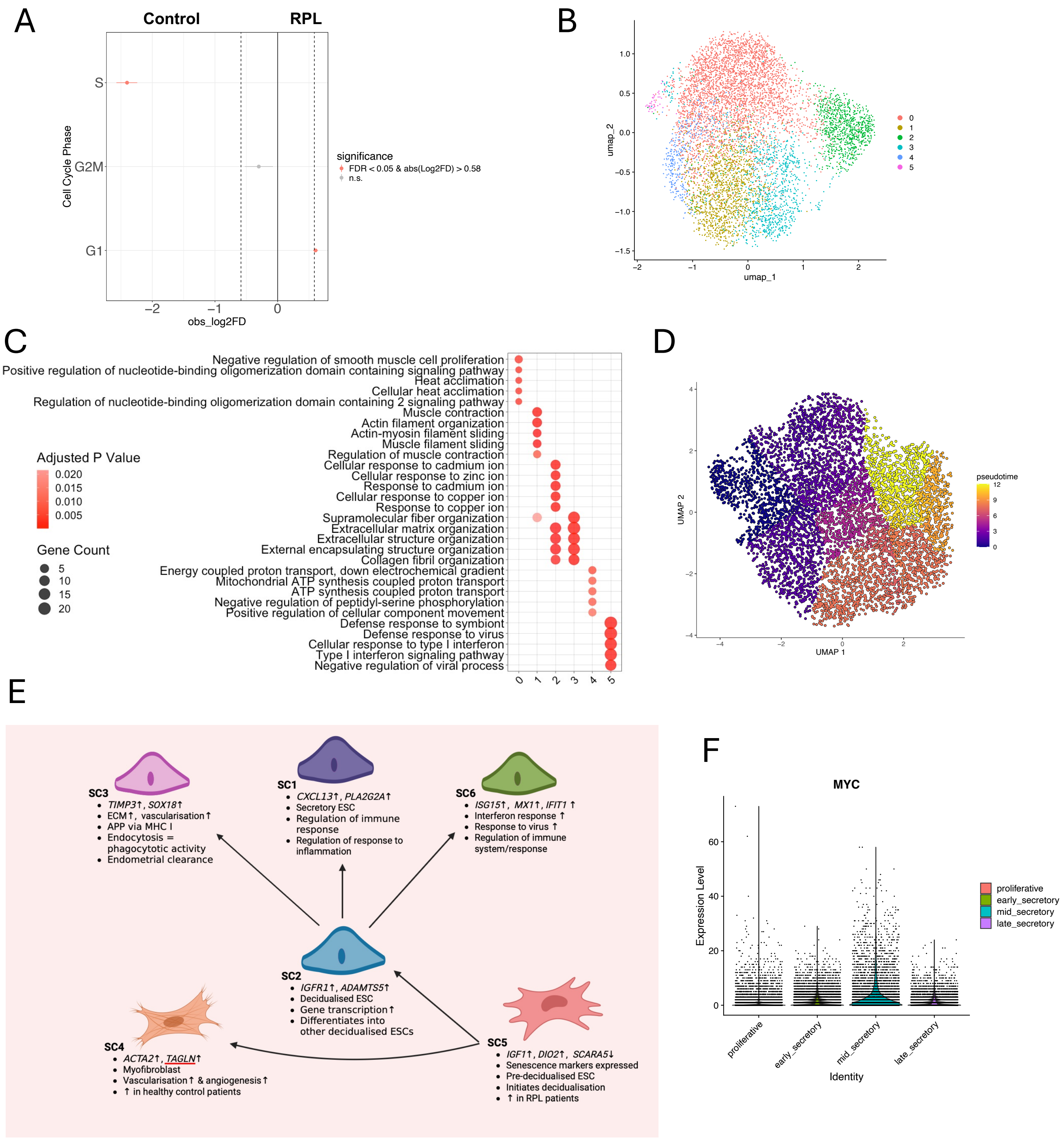


**Supplementary Figure S3 Endometrial stromal cells in the secretory phase. A)** ScProportionTest for cell cycle phase between control and RPL samples. **B)** UMAP of pseudotime scores from trajectory analysis using Monocle3. **C)** UMAP of clustered endometrial stromal cells (EnSCs) of the secretory phase in validation cohort. **D)** Dotplot showing top 5 GOBP pathways based on the significantly upregulated genes in each EnSC in the validation cohorts. **E)** Proposed EnSC subtypes in our cohort. **F)** *MYC* expression in validation cohorts by stage.
