## Supplementary Figure 4 for "Changes in the cellular composition of the endometrium during the implantation window are associated with recurrent pregnancy loss"

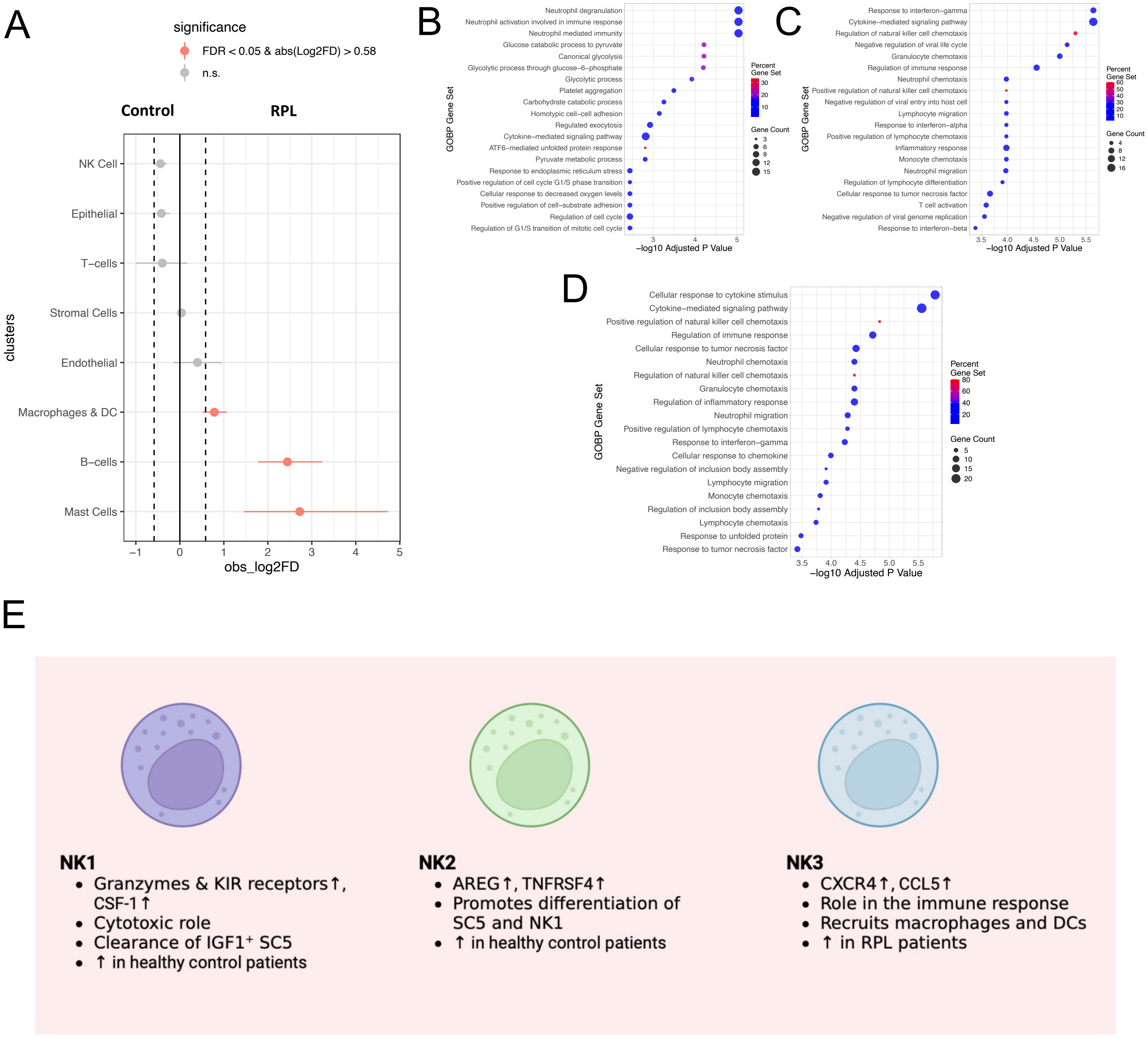


**Supplementary Figure S4 Natural killer cell populations in the endometrium. A)** Forest plot of differential proportions of major immune cell types between control and RPL groups. Top 20 GOBP pathways based on upregulated genes in **B)** NK1, **C)** NK2, and **D)** NK3. **E)** Summary of the three natural killer cell populations identified in our cohort.
