## Supplementary Figure 5 for "Changes in the cellular composition of the endometrium during the implantation window are associated with recurrent pregnancy loss"

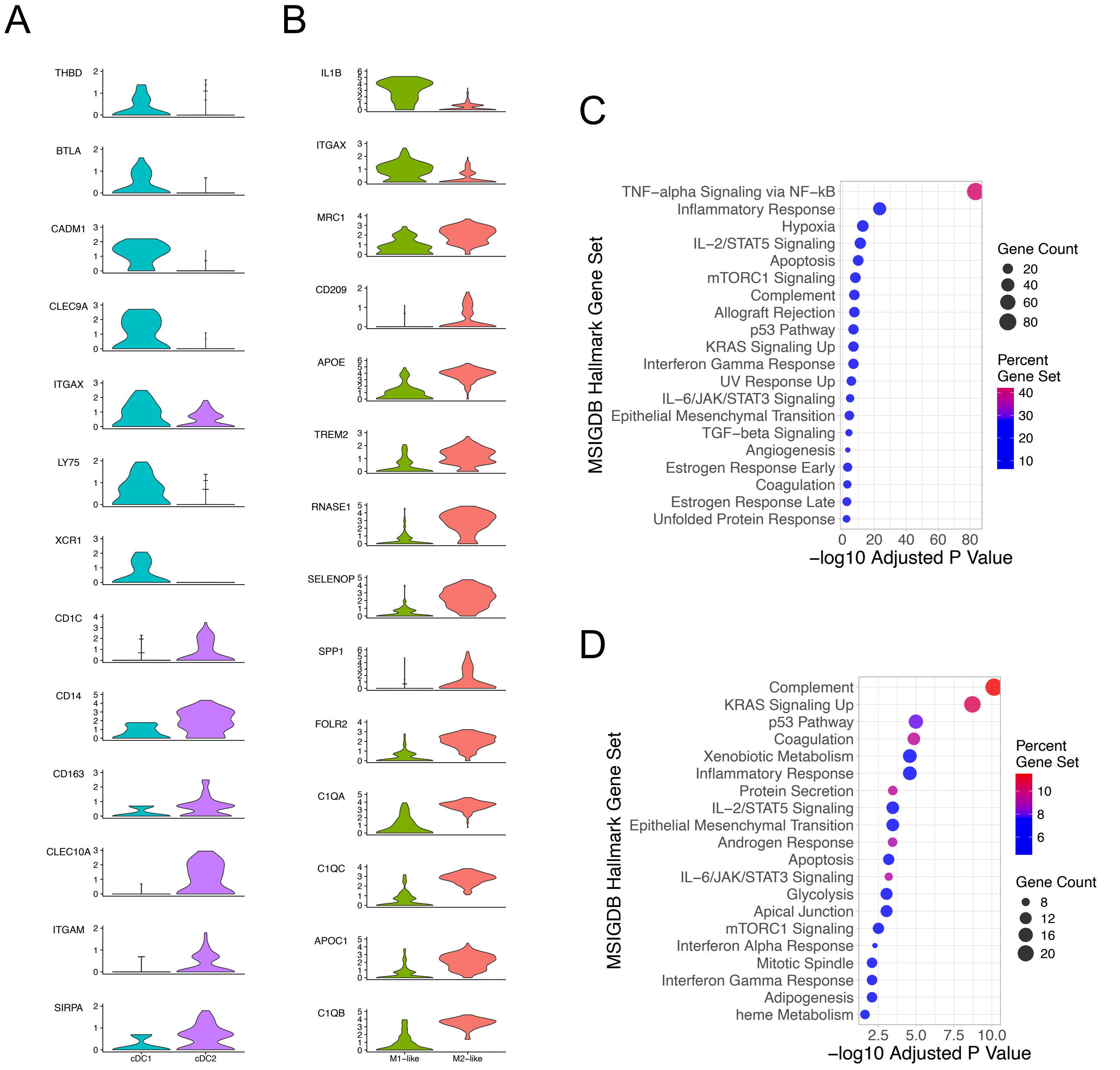


**Supplementary Figure S5 Macrophage and dendritic cell subpopulations in the secretory endometrium. A)** Violin plots of marker gene expression in M1-like and M2-like macrophage populations. Top 20 hallmark pathways based on the most upregulated genes in **B)** M1-like macrophages, and **C)** M2-like macrophages. **D)** Violin plots of marker gene expression in cDC1 and cDC2 dendritic cell populations. Top 20 GOBP pathways based on the most upregulated genes in **E)** cDC1, and **F)** cDC2 dendritic cells.
