## Supplementary Figure 6 for "Changes in the cellular composition of the endometrium during the implantation window are associated with recurrent pregnancy loss"

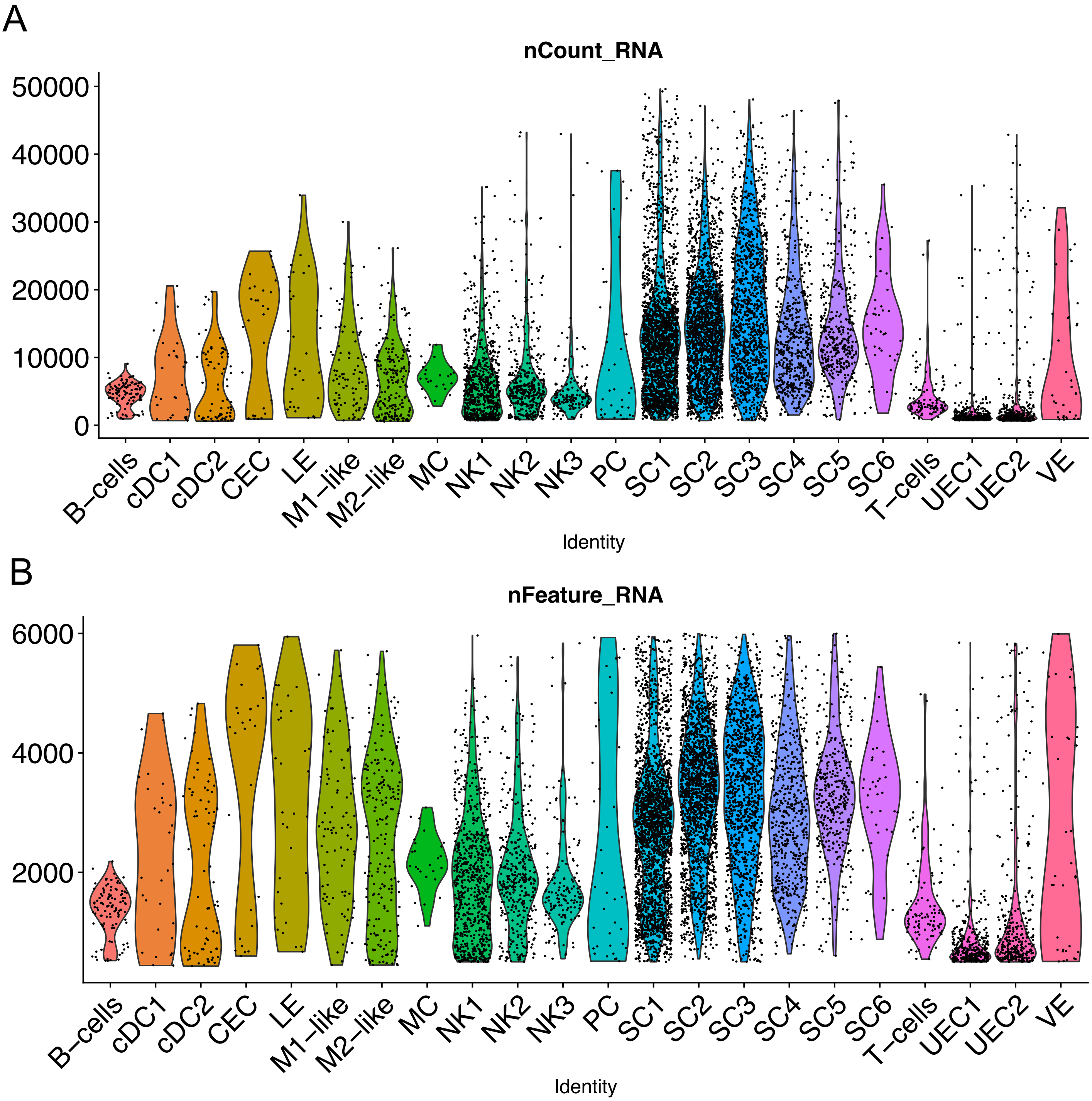


**Supplementary Figure S6 Gene expression is lower in unciliated epithelial cells.** Violin plots showing the number of reads **(A)**, and number of genes **(B)** expressed in each cell type.
