## Supplementary Figure 7 for "Changes in the cellular composition of the endometrium during the implantation window are associated with recurrent pregnancy loss"

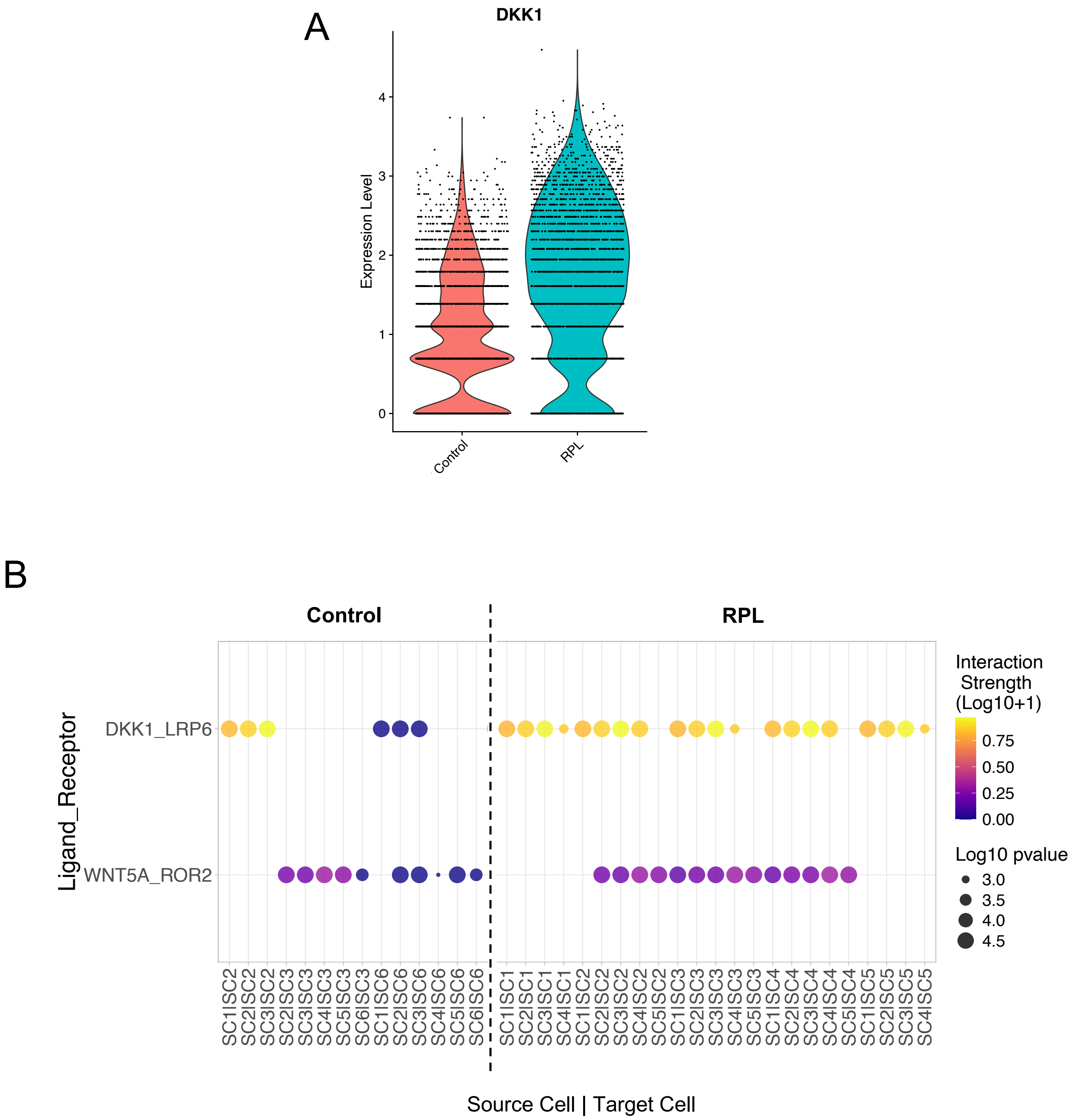


**Supplementary Figure S7 Wnt pathway signalling gene expression in the endometrium. A)** Violin plot of DKK1 expression in control and RPL groups. **B)** Dotplot of selected gene interactions associated with Wnt signaling In control and RPL groups.
